# Ubiquitination occurs in the mitochondrial matrix by eclipsed targeted components of the ubiquitin machinery

**DOI:** 10.1101/2022.04.21.488890

**Authors:** Yu Zhang, Ofri Karmon, Das Koyeli, Maayan Mark, Norbert Lehming, Ophry Pines

**Affiliations:** CREATE-NUS-HUJ Program and Department of Microbiology and Immunology, Yong Loo Lin School of Medicine, National University of Singapore, Singapore; Department of Molecular Genetics and Microbiology, IMRIC, Faculty of Medicine, Hebrew University, Jerusalem, Israel

## Abstract

Ubiquitination is a critical type of post translational modification in eukaryotic cells. It is involved in regulating nearly all cellular processes in the cytosol and nucleus. Mitochondria, known as the metabolism heart of the cell, are organelles that evolved from bacteria. Using the subcellular compartment-dependent α-complementation, we detect multiple components of ubiquitination machinery as being eclipsed distributed to yeast mitochondria. Subsequently, the results with respect to MTS (mitochondrial targeting sequence) targeted HA-tagged ubiquitin demonstrate that certain ubiquitination events specifically occur in yeast mitochondria and are independent of proteasome activity in the cytosol/nucleus. Importantly, we show that the E2 Rad6 affects the pattern of protein ubiquitination in mitochondria and provides an *in vivo* assay for its activity in the matrix of the organelle. This study shows that ubiquitination occurs in the mitochondrial matrix by eclipsed targeted components of the ubiquitin machinery, providing a new perspective of mitochondrial and ubiquitination research.

## INTRODUCTION

Ubiquitination is a type of eukaryote-specific protein modification in which the highly conserved 76 amino acid ubiquitin is covalently conjugated to a substrate protein. The process of ubiquitination is typically accomplished via a E1-E2-E3 enzymatic cascade involving ubiquitin-activating enzymes (E1), ubiquitin-conjugating enzymes (E2), and ubiquitin ligases (E3) [2]. Ubiquitination generally directs proteins to be degraded by the proteasome or lysosome. However, many ubiquitin modifications have non-proteolytic consequences, such as altered protein activity, cellular localization, and affinity to binding partners [3]. Ubiquitination is a reversible process, in which the deubiquitinating enzymes (DUBs) can reverse the process by removing ubiquitin from substrate proteins [4].

Mitochondria are vital eukaryotic organelles [5], hosting the tricarboxylic acid (TCA) cycle and oxidative phosphorylation to produce most of the cellular ATP. Mitochondria participate in critical cellular tasks such as amino acid and lipid metabolism, Fe-S clusters formation and heme biosynthesis, apoptosis and calcium signalling [6]. Disfunction of mitochondria is involved in aging, cancer, and pathological conditions/diseases of the nervous system, muscles, heart, and endocrine systems [7]. The involvement of ubiquitination in regulation and/or degradation of mitochondrial proteins has been previously reported. Early studies on this aspect were restricted to mitochondrial outer membrane (MOM) proteins due to the accessibility of these MOM proteins to the ubiquitination machinery in the cytosol. A number of studies in recent years revealed that the intramitochondrial proteins can be degraded through ubiquitination proteasome systems, but again the ubiquitination process for these proteins is reported to take place in the cytosol or in the space peripheral to mitochondria [8–10].

A version of the α-complementation assay was developed in our lab for in vivo detection of proteins distributed between mitochondria, especially mitochondrial matrix, and other cellular compartments in the model organism, budding yeast (*S. cerevisiae*) [11, 12]. Detection is possible even under low abundances of one of the compartments (termed “eclipsed distribution”) [13, 14]. Interestingly, in our recent screen for novel eclipsed/dual targeted mitochondrial proteins using the α- complementation assay (manuscript in preparation), initially, several components of ubiquitination system were detected in mitochondria. In this regard, there are three studies from two groups regarding ubiquitination of inner mitochondrial proteins. A bioinformatic study of the ubiquitinated proteome of human and yeast cells by the Ciechanover’s group revealed the existence of ubiquitinated mitochondrial proteins, including exclusive matrix proteins [15]. Subsequently, a biochemical study with isolated yeast mitochondrial samples by the same group, identified ubiquitinated mitochondrial matrix proteins [16]. Bénard’s group claimed that ubiquitination in mitochondria occurs with the detection of conjugates of mitochondrially directed HA-ubiquitin in the mitochondrial compartment of human HeLa cells [1]. Nevertheless, how the inner mitochondrial proteins undergo ubiquitination, where this occurs (in the mitochondrial matrix or intermembrane space, or in the cytosol), and by which components is still unclear. Our initial α-complementation results therefore raised the possibility that components of the ubiquitination system are imported and active in the *S. cerevisiae* mitochondrial matrix.

In this study, apart from the several E3s and a DUB initially detected in our screen, additional components of ubiquitination machinery of *S. cerevisiae* were detected in mitochondria by the α-complementation assay. Our biochemical studies employing MTS directed HA-Ubi (HA-tagged ubiquitin), indicate that ubiquitination occurs in mitochondria, and probably has a non-proteolytic function in *S. cerevisiae*. Furthermore, we show that the Rad6 E2 enzyme is eclipsed targeted to the *S. cerevisiae* mitochondria, which in turn is active locally (matrix) in protein ubiquitination as reflected by detection of an effect on the pattern of this modification in the organelle. Taken together, this study provides a new perspective regarding mitochondrial and ubiquitination research.

## RESULTS

### The α-complementation assay indicates targeting of components of ubiquitination system to mitochondria of *S. cerevisiae*

The α-complementation assay is based on the reconstitution of enzymatic activity of β-galactosidase when the two fragments (α - 77 amino acids and ω - 993 amino acids) of the enzyme co-localize in the same compartment [12]. Specifically, the short α fragment of β-galactosidase is fused to the C terminus of the protein of interest and is co-expressed with the compartmentalized ω fragment in *S. cerevisiae*. If the fusion protein of interest co-localizes with the ω fragment in the same compartment within the cell, the reconstitution of β-galactosidase activity causes formation of blue colonies on agar plates supplemented with X-Gal (the substrate of β-galactosidase). Otherwise, the colour of the colonies will be white (Figure 1A). This approach is sensitive for detection of eclipsed distribution of a protein, i.e., the presence of significantly minute amounts of a protein in a second cellular location [13].

**Figure 1:**
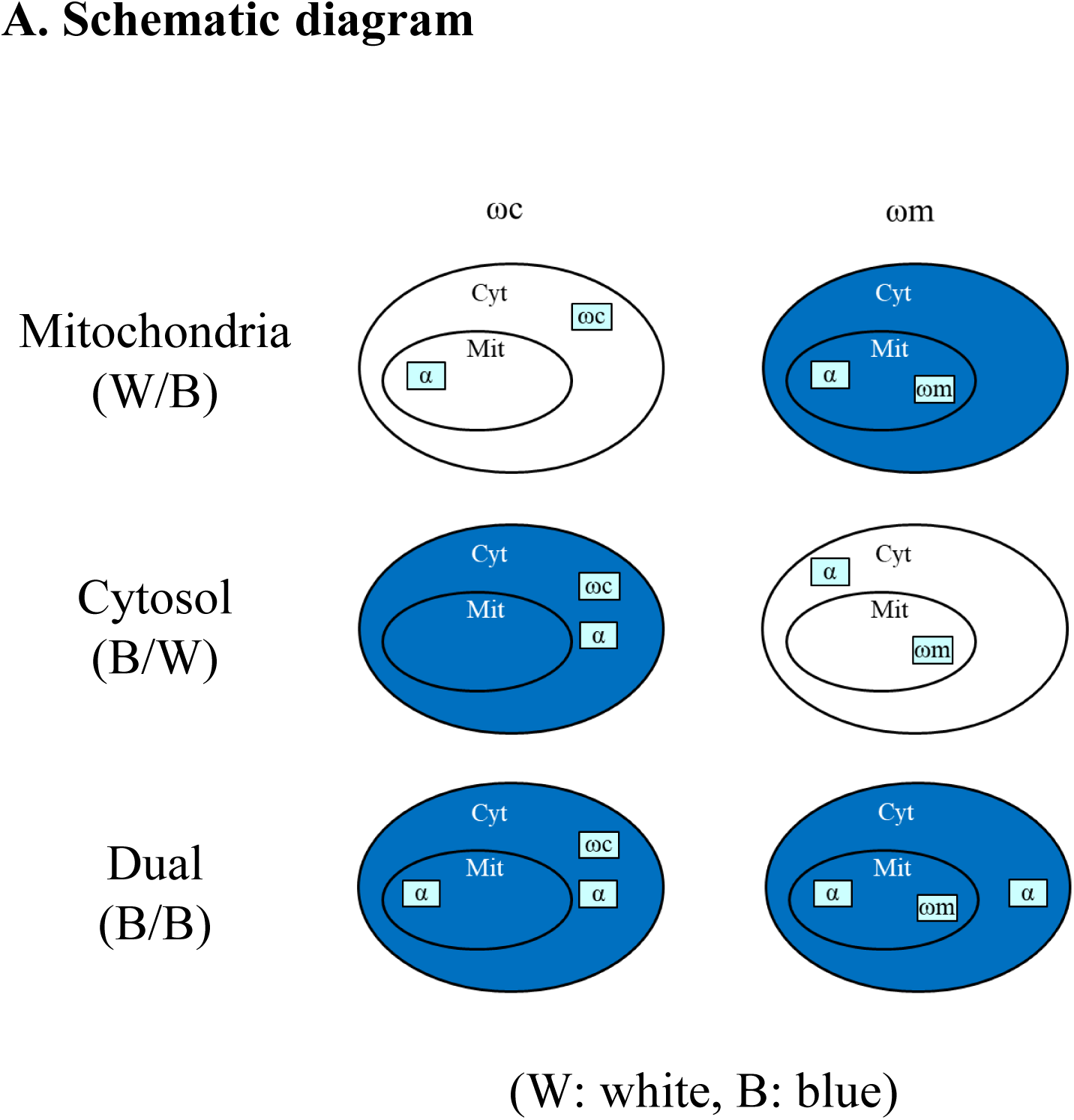
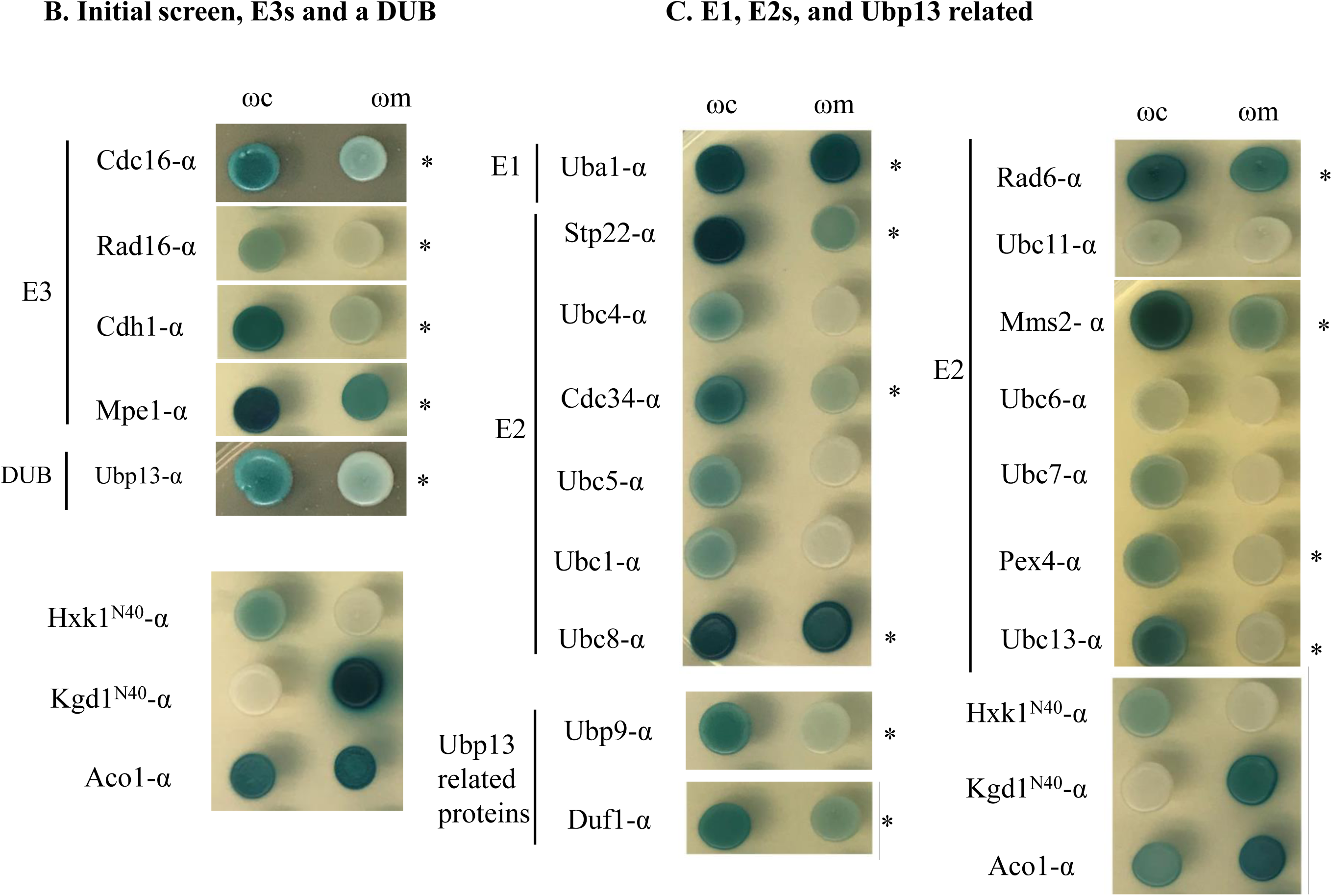
The α-complementation assay identifies candidate components of the ubiquitination machinery in mitochondria. **(A) Schematic diagram of the α-complementation assay.** Proteins of interest were fused with the α-fragment of β-galactosidase and expressed in cells with the ω- fragment present either in cytosol (Cyt) or mitochondria (Mit). If α and ω fragments colocalize in the same compartment, the cell colony colour will be blue, and if the two fragments of β-galactosidase are present in separate compartments, the cell colony colour will be white. **(B) Initial screen, E3s and a DUB are targeted to mitochondria.** Four E3s including Cdc16, Rad16, Cdh1, MPE1, and a DUB, Upb13, were detected in mitochondria in the initial α-complementation screen of hundreds of proteins not known to be mitochondrial. N terminal 40 amino acids of Hxk1 (Hxk1^N40^) and Kgd1 (Kgd1^N40^) are cytosolic and mitochondrial markers, respectively, and Aco1 is a dual targeted marker. **(C) Mitochondrial targeting of E1, E2s and Ubp13 related proteins.** Uba1, Stp22, Cdc34, Ubc8, Rad6, Mms2, Pex4, Ubc13, Ubp9, and Duf1 are mitochondrial positive by the α-complementation assay. * indicates a blue colour with ωm.

Recently we performed a screen for novel dual targeted *S. cerevisiae* mitochondrial proteins by using the α-complementation assay (manuscript in preparation). Several α-fusions of the components of the ubiquitination system were detected both in mitochondria and the cytosol, including four E3 enzymes; Cdc16, Rad16, Cdh1 and Mpe1, and a DUB, Ubp13 (Figure 1B). Accordingly, we raised the possibility that these and other components of the ubiquitination system may be imported and active in protein ubiquitination in mitochondria.

We next investigated the mitochondrial targeting of ubiquitination cascade enzymes, functioning upstream of E3 ligases, using the α-complementation assay. E2s including Cdc34 (Ubc3), Gid3 (Ubc8), Rad6 (Ubc2), Pex4 (Ubc10), Ubc13, Stp22 [15], and Mms2, (the ubiquitin E2 variant, UEV, protein known to act in concert with Ubc13), exhibited positive blue when co-expressed with the ω fragment either in the cytosol (ωc) or mitochondria (ωm). The same is true for the single E1, Uba1. This suggested mitochondrial targeting of the E1 and 7 E2 (related) proteins (Figure 1C). In addition, the paralog of Ubp13, Upb9, and the activator of the two DUBs, Duf1, are suggested to also be targeted to mitochondria as well (Figure 1C).

To further explore the possibility of ubiquitination in *S. cerevisiae* mitochondria, we performed subcellular fractionation and examined ubiquitin conjugates in mitochondrial fractions by western blotting (WB). Separation between mitochondrial and cytosolic fractions was achieved by standard procedures. Briefly, homogenized yeast spheroplasts were subjected to low-speed centrifugation to remove debris and then to high-speed centrifugation (10000x g) to separate mitochondria. The 10000x g cleared lysate and the pellet of isolated mitochondria after being resuspended and homogenized, are defined as cytosolic and mitochondrial fractions, respectively (Figure 2, top panel). Hsp60 and Hxk1 were employed as the mitochondrial and cytosolic reference markers respectively (Figure 2, middle panel). Since mitochondria contribute a small percentage to total cellular protein mass [17], a fifteen-fold (15X) enriched mitochondrial fraction in addition to non-enriched (1X) mitochondrial fraction was subjected to WB using anti-ubiquitin antibodies (Figure 2, top panel, M^15x^). Figure 2, top panel, shows that the ubiquitin conjugates signal can be clearly detected in the cytosolic and in the 15X enriched mitochondrial fraction. Not surprisingly, the ubiquitin conjugates signal of the 1X mitochondrial fraction is weak due to the very low level of protein mass (compare lanes of the bottom panel of Figure 2). The fractions in Figure 2 were prepared from galactose grown cells since further on in this study the galactose induced promoter is used for expression of proteins of interest from expression plasmids. Nevertheless, similar corresponding results were obtained with cells grown on glucose (Supplementary Figure S1). These results further validate the possibility of protein ubiquitination in mitochondria.

**Figure 2:**
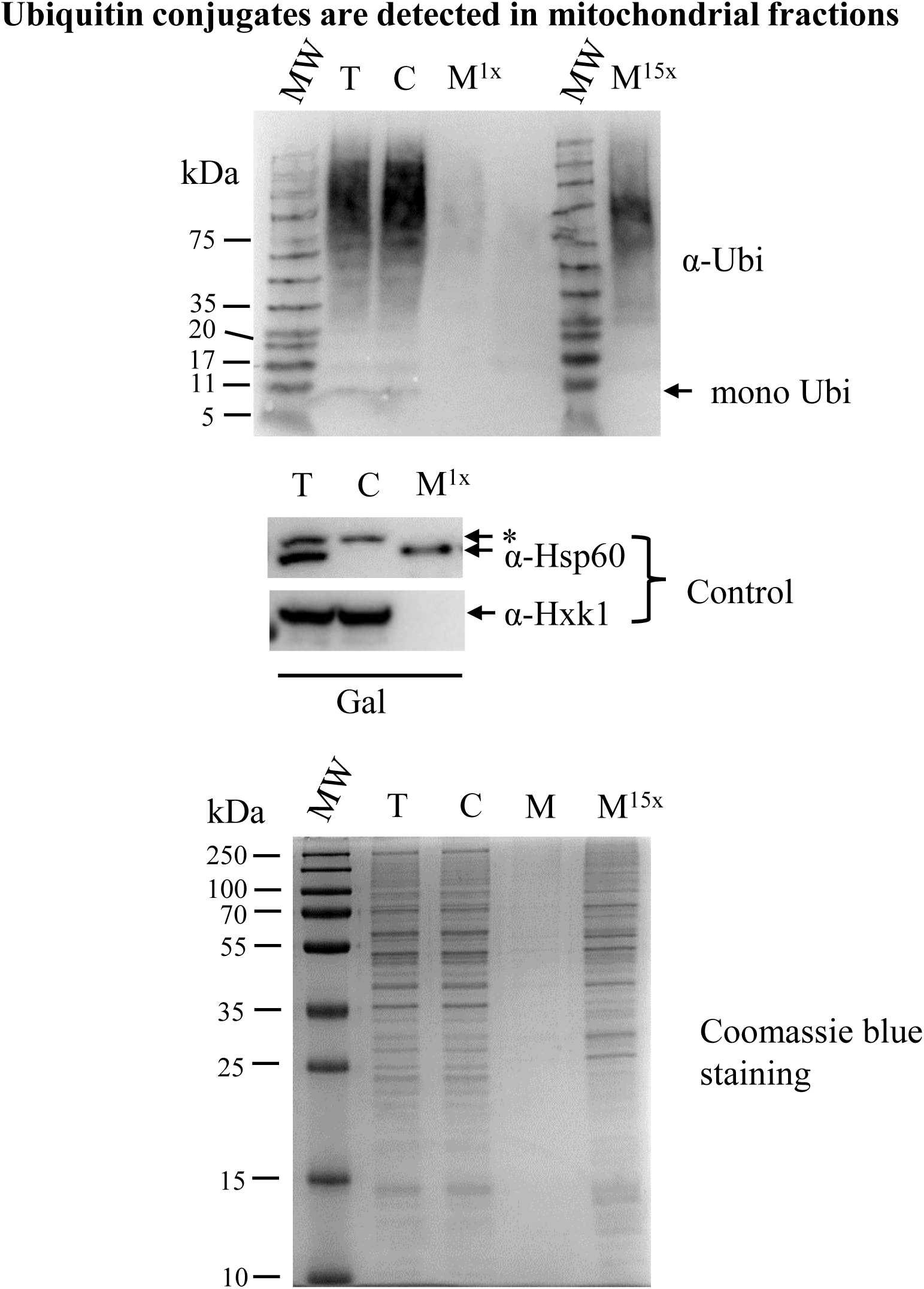
Ubiquitin conjugates are detected in mitochondrial fractions. **Top Panel: Subcellular fractions probed for ubiquitin.** Subcellular fractions were prepared from cells grown to logarithmic phase in galactose minimal medium. Mitochondrial fractions (M) corresponding to 1Χ and 15Χ cells, total (T) and cytosolic (C) fractions corresponding to 1Χ cells were subjected to western blot employing anti-ubiquitin (α-Ubi) antibodies. **Middle Panel:** T, C, and M fractions were probed with control antibodies against Hsp60 and Hxk1, which are mitochondrial and cytosolic markers, respectively. *, the upper bands detected in T and C fractions are non-specific. **Bottom Panel:** Subcellular fractions were subjected to 12.5% SDS-PAGE and Coomassie Blue staining. MW in the figure and figures elsewhere in the present study refers to molecular weight.

### Targeting HA-tagged ubiquitin to *S. cerevisiae* mitochondria

#### Conjugates of MTS directed HA-Ubi are detected only in mitochondrial fractions

To specifically follow ubiquitination in mitochondria we expressed two HA-tagged versions of ubiquitin (HA-Ubi) constructs, harbouring the MTS of Su9 (preSu9-HA-Ubi) and fumarase (preFum1-HA-Ubi), respectively, under the yeast *MET25* promoter. In addition, we expressed HA-Ubi lacking an MTS as a control (Figure 3A). If ubiquitination occurs in mitochondria, the conjugates of the MTS directed HA-Ubi are expected to be specifically detected in mitochondria by HA antibody. Indeed, signals for ubiquitin (HA-tagged) conjugates were clearly detected in mitochondrial enriched fractions, rather than in the cytosolic fractions of cells expressing either preSu9-HA-Ubi or preFum1-HA-Ubi (Fig 3B). In contrast, for cells expressing the control HA-Ubi (without an MTS), conjugate signals were detected in both mitochondrial and cytosolic fractions, and no conjugate signals were detected, at all, in cells harbouring the empty plasmid (no HA-Ubi) (Fig 3B). The Su9 MTS (preSu9) is a well-established mitochondrial matrix targeting signal [18–20] and therefore, preSu9-HA-Ubi was employed for the rest of this study. Taken together these observations indicate the occurrence of protein ubiquitination in mitochondria.

**Figure 3:**
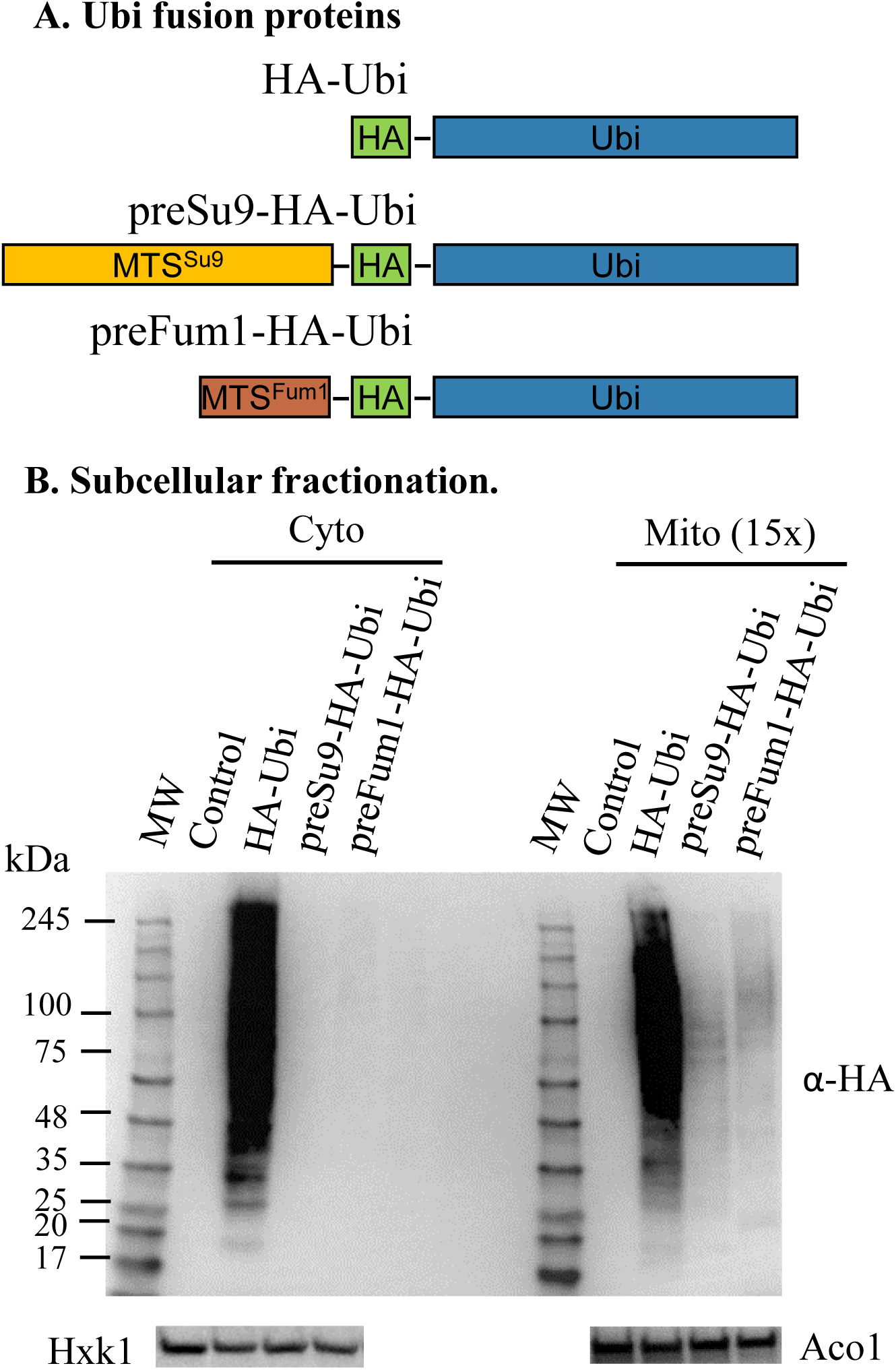
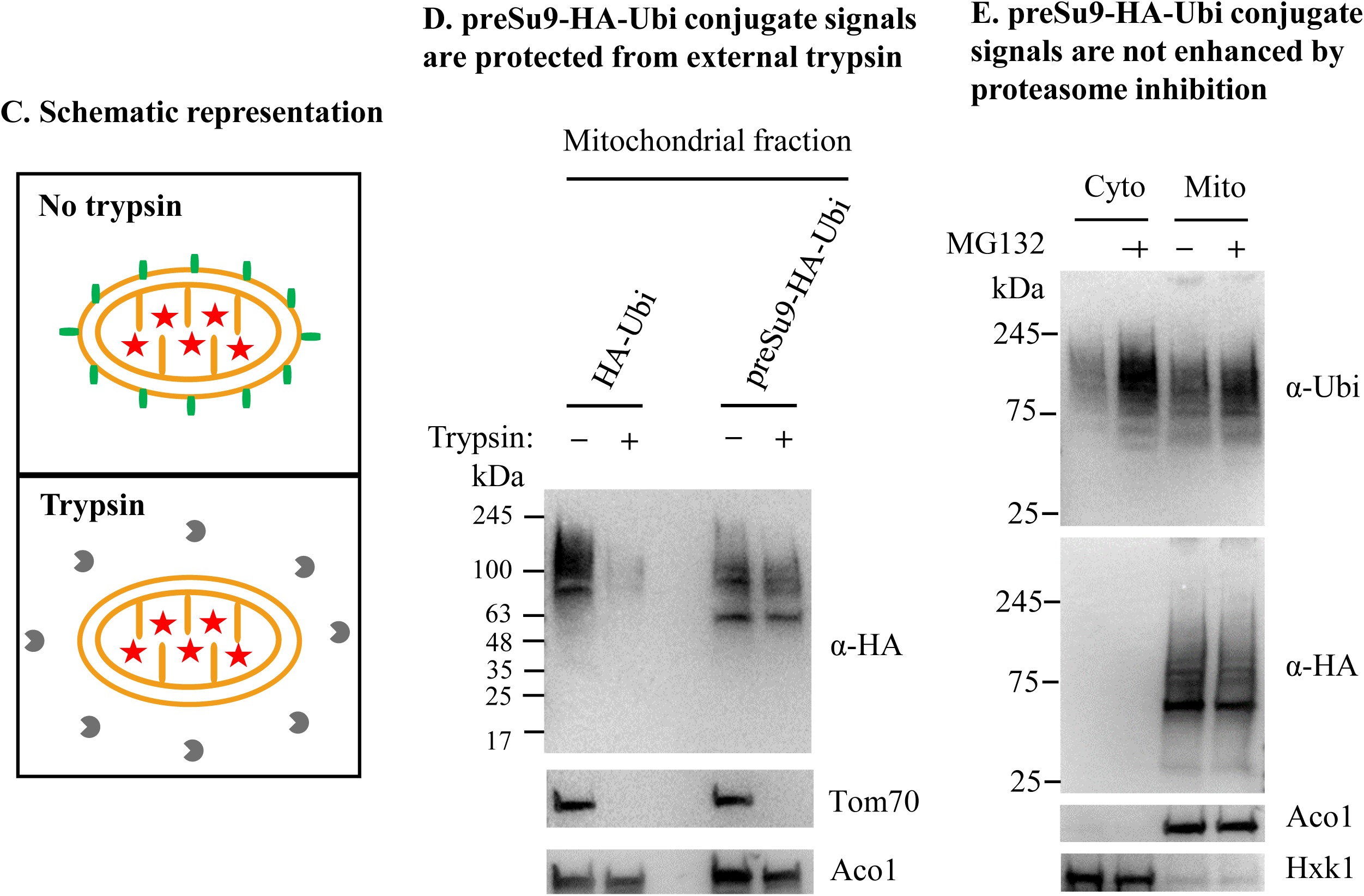
Conjugates of mitochondrially directed MTS-HA-Ubi are detected in mitochondria. **(A) Schematic representation of ubiquitin fusion protein constructs.** Three ubiquitin fusion proteins were constructed, including an N-terminal HA-tagged ubiquitin (HA-Ubi), and two mitochondrially directed HA-Ubi by adding the presequence of subunit 9 of F0-ATPase of *Neurospora crassa* (1-69 aa) or Fum1 (1-27 aa) in front of HA-Ubi, which are respectively named preSu9-HA-Ubi and preFum1-HA-Ubi. **(B) Detection of HA-Ubi in mitochondria by western blot.** Subcellular fractions were prepared from glucose grown BY4742 *Δpdr5* cells expressing HA-Ubi, preSu9- HA-Ubi, preFum1-HA-Ubi or empty plasmid (Control). Mitochondrial fractions (Mito) corresponding to 15Χ cells, and cytosolic fractions (Cyto) corresponding to 1Χ cells were subjected to western blot for HA-marked ubiquitin conjugates detection using *α*-HA antibodies. Hxk1 and Aco1 antibodies served as loading and fractionation controls. **(C) Schematic representation on the trypsin treatment of isolated mitochondria.** The mitochondrial membranes (), an outer membrane protein exposed to trypsin (), a matrix protein protected inside mitochondria () and trypsin () are indicated. **(D) preSu9-HA-Ubi conjugate signals are protected from external trypsin.** Mitochondrial fractions were prepared from cells expressing HA-Ubi or preSu9-HA- Ubi and mitochondrial fractions with and without trypsin treatment were subjected to western blot using *α*-HA antibodies. Tom70, a cytosol-facing mitochondrial outer membrane protein, was used as a marker exposed to the trypsin treatment, and Aco1 is a protected mitochondrial matrix marker. **(E) preSu9-HA-Ubi conjugates signals are not enhanced by proteasome inhibition treatment.** Cytosolic (Cyto) and mitochondrial (Mito) subcellular fractions prepared from preSu9-HA-Ubi expressing cells in the presence or absence of MG132 treatment were subjected to western blot. Detection of general and HA- tagged ubiquitin conjugates signals employed α-Ubi and α-HA antibodies, respectively. Cytosolic marker Hxk1 and mitochondrial marker Aco1 are indicated.

#### Conjugate signals of preSu9-HA-Ubi in isolated mitochondria are resistant to external trypsin treatment

Unlike mitochondrial peripheral proteins and mitochondrial outer membrane (MOM) proteins facing the cytosol, preSu9-HA-Ubi conjugates from isolated mitochondria are assumed to be localized in the matrix and resistant to trypsin treatment of whole mitochondria (Figure 3C). Indeed, the preSu9-HA-Ubi conjugates signals of the isolated mitochondria with and without trypsin treatment exhibit a similar pattern and signal strength (Figure 3D, top panel, two right lanes). In contrast, as expected, HA-Ubi modified proteins which are mainly present outside mitochondria are sensitive to trypsin (Figure 3D, top panel, two left lanes). In this experiment the mitochondrial MOM protein, Tom70, was used as an outer membrane marker protein, exposed to the cytosol, which can be accessed by trypsin in vitro. Accordingly, the Tom70 control was completely undetectable after trypsin treatment of isolated mitochondria while the matrix protein Aco1 was protected from trypsin digestion (Figure 3D, middle and bottom panels, respectively). The quality of the subcellular fractionation is shown in Figure S2. These data together indicate that the mitochondrially directed ubiquitin conjugates are protected inside mitochondria and thereby support the notion of protein ubiquitination in mitochondria.

#### preSu9-HA-Ubi conjugates levels in mitochondria are not elevated by proteasome inhibition

Ubiquitinated proteins in the cytosol or nucleus are in most cases degraded in the proteasome. In fact, inhibition of the proteasome by chemical inhibitors such as MG132 is known to significantly increase the level of ubiquitinated proteins in the cell [21]. In order to test whether the ubiquitination we observe is related to proteasome degradation or other pathways, we blocked proteasome activity using MG132. While MG132 treatment increased the ubiquitination level in the cytosolic fraction (Figure 3E, top panel, α-Ubi, two left lanes), preSu9-HA-Ubi conjugates levels of the mitochondrial fraction was not elevated (Figure 3E, second panel, α-HA, compare the two right lanes). This suggests that preSu9-HA-Ubi conjugate signals in mitochondria are not exposed to the cytosolic proteasome and that the preSu9-HA-Ubi ubiquitinated proteins in mitochondria may have a different fate. In this regard, it is important to state that our α-complementation screen only detected candidate components of the ubiquitination system but none of the proteasome (manuscript in preparation). These results demonstrate ubiquitination of proteins in mitochondria. It is not surprising that the ubiquitination level of the mitochondrial fraction detected by α-Ubi (top panel, two right lanes) increased following MG132 treatment, since the ubiquitin antibody recognizes outer membrane proteins and proteins associated with the mitochondrial outer surface, which may be proteasome substrates.

#### Immunoprecipitation (IP) of preSu9-HA-Ubi conjugates is enriched in mitochondrial matrix proteins

If conjugation of mitochondrially directed ubiquitin occurs in mitochondria, the substrates of preSu9-HA-Ubi are expected to be matrix proteins or inner mitochondrial membrane proteins facing the matrix. This is because the preSu9p is a well-established strong mitochondrial matrix targeting signal as mentioned earlier. Therefore, the protein composition immunoprecipitated by anti-HA magnetic beads from mitochondrial fractions of cells expressing preSu9-HA-Ubi, was examined.

Equal OD_600_ units of cells expressing preSu9-HA-Ubi or HA-Ubi, or harbouring an empty plasmid (Control) were used to prepare mitochondrial subcellular fractions, which were then subjected to IP with α-HA. The proteins in the immunoprecipitated (IPed) samples after trypsinization were subsequently identified by mass spectrometry (Figure 4A). Mitochondria were efficiently isolated from each type of cells, as the mitochondrial marker Tom70 was only detected in mitochondrial fractions (Figure S3A). Figure S3B shows that HA-tagged ubiquitin-protein conjugates were detected for preSu9-HA-Ubi and HA-Ubi IP but not for Control IP. The detailed list of the IPed proteins identified by MS can be found in Table S1. Table 1 shows the number of proteins identified in each of the two parallel IPs (preSu9-HA-Ubi and HA-Ubi IPs), after subtracting proteins detected in the Control IP; 131 and 557 proteins are in the list of proteins IPed from mitochondrial lysates of preSu9-HA-Ubi and HA-Ubi expressing cells, respectively. Two thorough studies on the mitochondrial proteome of yeast were published by Vogtle et al. (2017) and Morgenstern et al. (2017) respectively, both of which examined the protein contents of mitochondrial sub-compartments [22, 23]. These two studies are our reference when looking at the composition of the two groups (131 and 557 referred to above) of proteins (Table 1). Regarding preSu9-HA-Ubi IP, 109 out of the 131 (83.2%) are found in the mitochondrial proteome and among these 109 proteins, 78 (71.6%) are reported to be in the mitochondrial matrix (Vogtle et al. and Morgenstern et al). In contrast, for HA-Ubi IP, the fraction of proteins belonging to mitochondrial proteome is significantly much lower (48.3%, 269 out of the 557), indicating that the substantial surplus of HA-Ubi conjugates, IPed out, are probably externally exposed on mitochondria. This is consistent with the results of external trypsin treatment (Figure 3D). Out of the 269 mitochondrial proteins detected for HA-Ubi IP, 25.7% (69/269) are considered matrix proteins, which is dramatically lower than 71.6% for the preSu9-HA-Ubi IP (Table 1, Figure 4B). Thus, we conclude that immunoprecipitation of preSu9-HA-Ubi conjugates is highly enriched for proteins in mitochondrial matrix, supporting specific ubiquitination in the mitochondrial matrix.

**Figure 4:**
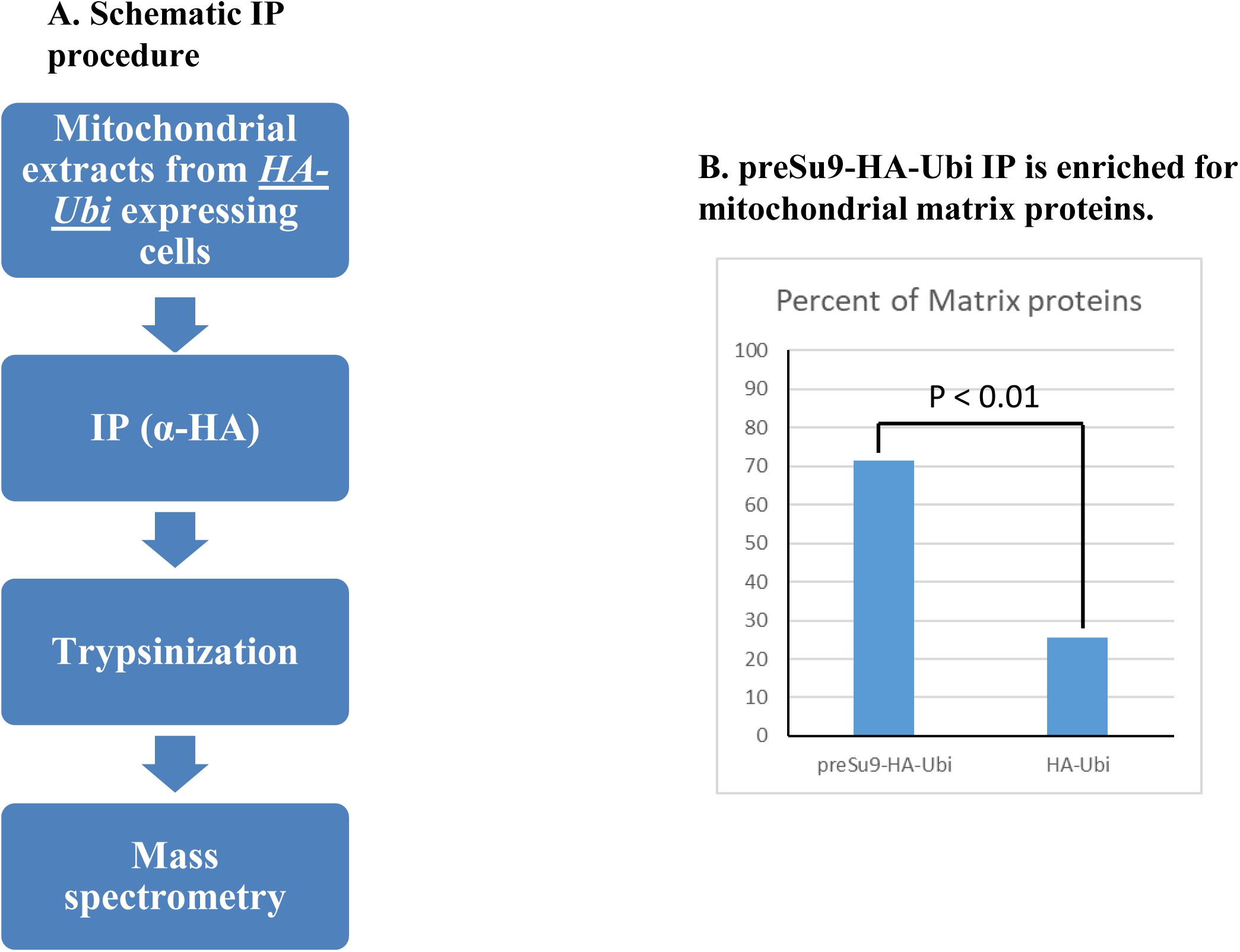
Immunoprecipitation (IP) of ubiquitinated HA containing conjugates. **(A) Schematic representation of the IP.** IPs were performed with anti-HA magnetic beads incubated with mitochondrial lysates of cells harboring empty vector or expressing preSu9-HA-Ubi or HA-Ubi. IPed samples after trypsinization were subjected to mass spectrometry (MS). **(B) Percent of matrix proteins detected by MS for preSu9-HA-Ubi and HA-Ubi IP:** 71.6% (78/109) and 25.7% (69/269) respectively. The numerator in each bracket is the number of matrix proteins detected, the denominator is the number of mitochondrial proteins detected (Table 1).

**Table 1:** IP of preSu9-HA-Ubi conjugates is enriched for mitochondrial matrix proteins.

| IP | No. of protein IDs<sup>a</sup> | No. of Mito proteins | % of Mito proteins<sup>b</sup> | No. of matrix proteins | % of matrix proteins<sup>c</sup> |
| --- | --- | --- | --- | --- | --- |
| HA-Ubi | 557 | 269 | 48.3% | 69 | 25.7% |
| preSu9-HA-Ubi | 131 | 109 | 83.2% | 78 | 71.6% |

<sup>a</sup> 423 protein IDs from cells not expressing HA (Control) were subtracted. <sup>b</sup> and <sup>c</sup>, Chi-squared test was used to determine that the observed frequency difference between the indicated two IPs was statistically significant. The two p values are both far less than 0.01.

### RAD6 deficiency causes an altered ubiquitination pattern in mitochondria

preSu9 MTS directed HA-Ubi (preSu9-HA-Ubi), allows detection of ubiquitination in mitochondria. We asked whether the components of ubiquitination machinery, which were shown to be targeted to mitochondria by α-complementation, have an effect on the detectable pattern of ubiquitination in mitochondria. Our initial strategy was to look at the pattern of preSu9-HA-Ubi ubiquitinated proteins in the mitochondria of cells in which the gene encoding a component of ubiquitination machinery was deleted (“KO’ed”). E3s generally confer the specificity of ubiquitination and in yeast there are approximately 80 E3s, while there is only one E1 and 11 E2s [24]. Therefore, we speculated that there would be a higher chance to detect an effect of an E1 or E2 on the pattern of ubiquitination in mitochondria. Among the E1 and E2 (related) proteins that are suggested to be targeted to mitochondria by the α-complementation assay (Fig 1B and C), five E2 (related) proteins including Mms2, Pex4, Rad6, Ubc8, and Ubc13 have a non-lethal KO phenotype. We decided to examine these five individual E2 KO strains.

Figure 5 shows that there were insignificant differences in the α-Ubi probed general patterns of ubiquitination, in the different strains. The mitochondrial ubiquitination pattern of α-HA was probed for preSu9-HA-Ubi conjugates. A major discovery for us, here, was the significant difference between mitochondrial fractions of RAD6 KO strain and wild-type (WT) and other KO strains. More specifically, the most pronounced difference is in the “major band” of preSu9-HA-Ubi conjugates signals, between 48 and 63 kDa (this is the major band referred to hereafter); this band is present in WT cells and is absent in cells with the RAD6 KO (Figure 5, top panel). Beyond the major band difference, there are other, less pronounced differences between the preSu9- HA-Ubi conjugates-signals of RAD6 KO cells versus WT cells.

**Figure 5:**
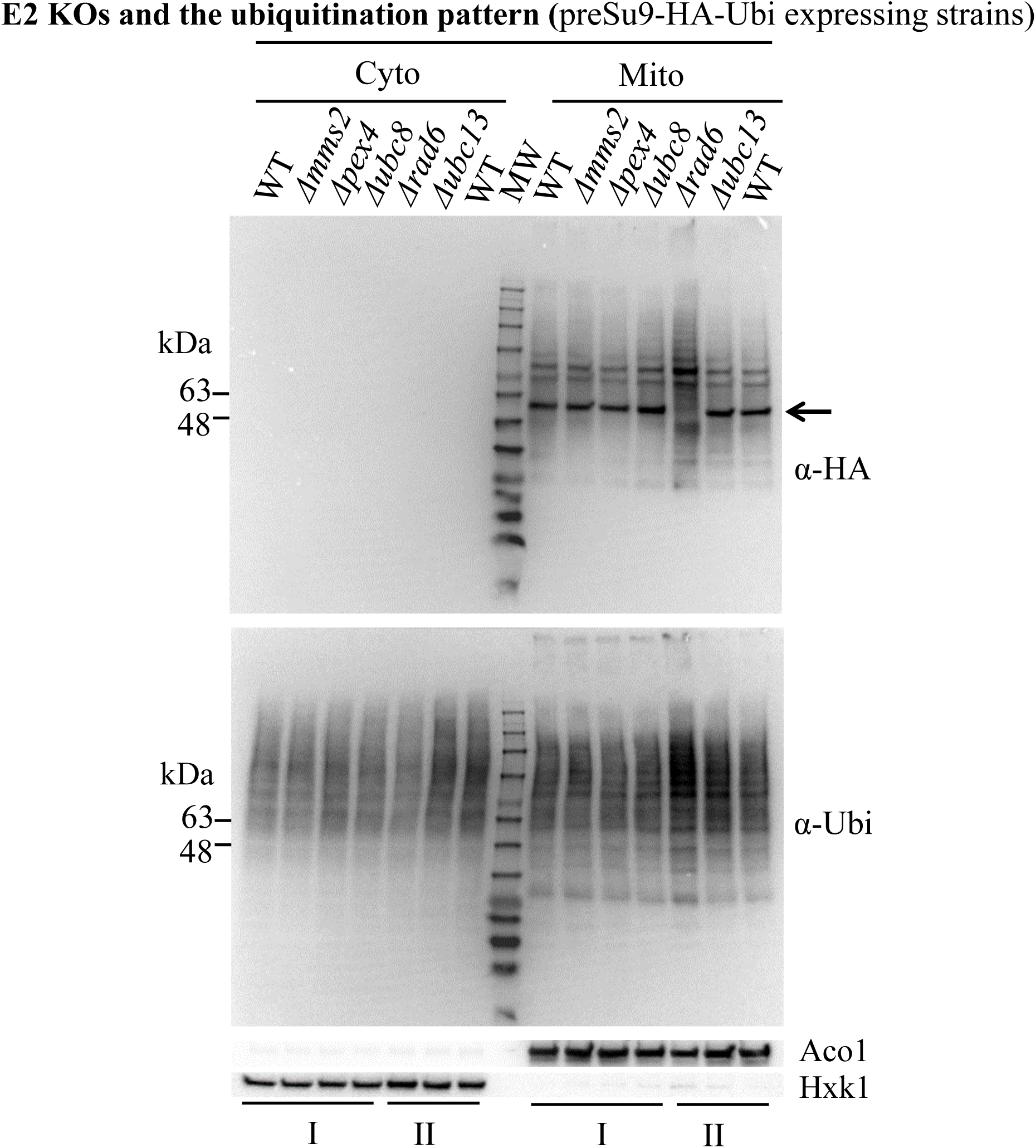
RAD6 deficiency causes an altered mitochondrial ubiquitination pattern. Effect of single E2 KO on the mitochondrial ubiquitination pattern. Wild-type (WT) or single E2 KO cells expressing preSu9-HA-Ubi were subjected to subcellular fractionation. Cytosolic (Cyto) and mitochondrial (Mito) subcellular fractions were probed for HA or ubiquitin (Ubi) by WB. Aco1 and Hxk1 respectively are mitochondrial and cytosolic markers. Line arrow indicates the migration position of the major band between 48 and 63 kDa. Samples prepared from two batches of subcellular fractionation are indicated with I and II.

We assumed that the major band is probably formed by ubiquitinated substrate(s) whose ubiquitination requires the activity of Rad6 in mitochondria and probably involves a Rad6 interacting E3. Rad6 is known to form a ubiquitin ligase complex with E3s in ubiquitinating substrate proteins in the nucleus and cytosol. The identified ubiquitin ligase complexes with Rad6 include RAD6-RAD18, BRE1-RAD6, UBR1- RAD6, RAD6-UBR2, and MUB1-RAD6-UBR2 complexes, which are required for a variety of cellular processes including post-replicative repair of damaged DNA, epigenetic transcriptional activation, modulating the formation of double-strand breaks during meiosis, DNA-damage checkpoint activation, telomeric gene silencing, N-end rule protein degradation, endoplasmic reticulum (ER)-associated protein degradation (ERAD), and proteasome homeostasis [25–30]. We tested whether any of the 5 complex proteins mentioned above, i.e., Rad18, Bre1, Ubr1, Ubr2, and Mub1, is required for the formation of the major band in mitochondria. We also tested the preSu9-HA-Ubi conjugates pattern in mitochondria in another 12 strains deficient in Ela1, Cos111, Cdh1, Rad16, Rav1, Hul5, Bul2, Rtt101, Mmd2, Asi3, Hrd1, and Nam1, respectively. These 12 proteins are the E3s which scored highest by the program MitoProt II in terms of containing an N-terminal MTS and/or testing positive for mitochondrial targeting in our α-complementation screen. Additionally, we also tested a strain deficient in E3, Dma1, found to be present in mitochondrial matrix [16]. Unfortunately, none of these 18 proteins appear to be specifically required for the normal ubiquitination pattern in mitochondria and thus are not exclusive Rad6 interacting E3s in mitochondria (Figure S4).

### Rad6 is active in mitochondria

According to these results we decided to focus on Rad6 and use its effect on the ubiquitination pattern in mitochondria as an assay for Rad6 function in mitochondria. First, we introduced C terminally α-tagged Rad6 (Rad6-α) into the RAD6 KO cells and found that the WT pattern of preSu9-HA-Ubi conjugates was restored (Figure 6A, top panel, third lane from the right). In addition, Rad6-α restored the normal growth of RAD6 KO cells (Figure 6B). No restoration of the ubiquitination pattern and growth rate was found using a construct of α-tagged Ubc8 (Ubc8-α), an E2 suggested to be targeted to mitochondria in our α-complementation assay (Figure 6A and B). Taken together, these results indicate a clear role of Rad6 in affecting the ubiquitination of proteins in mitochondria and for maintaining normal cell growth.

**Figure 6:**
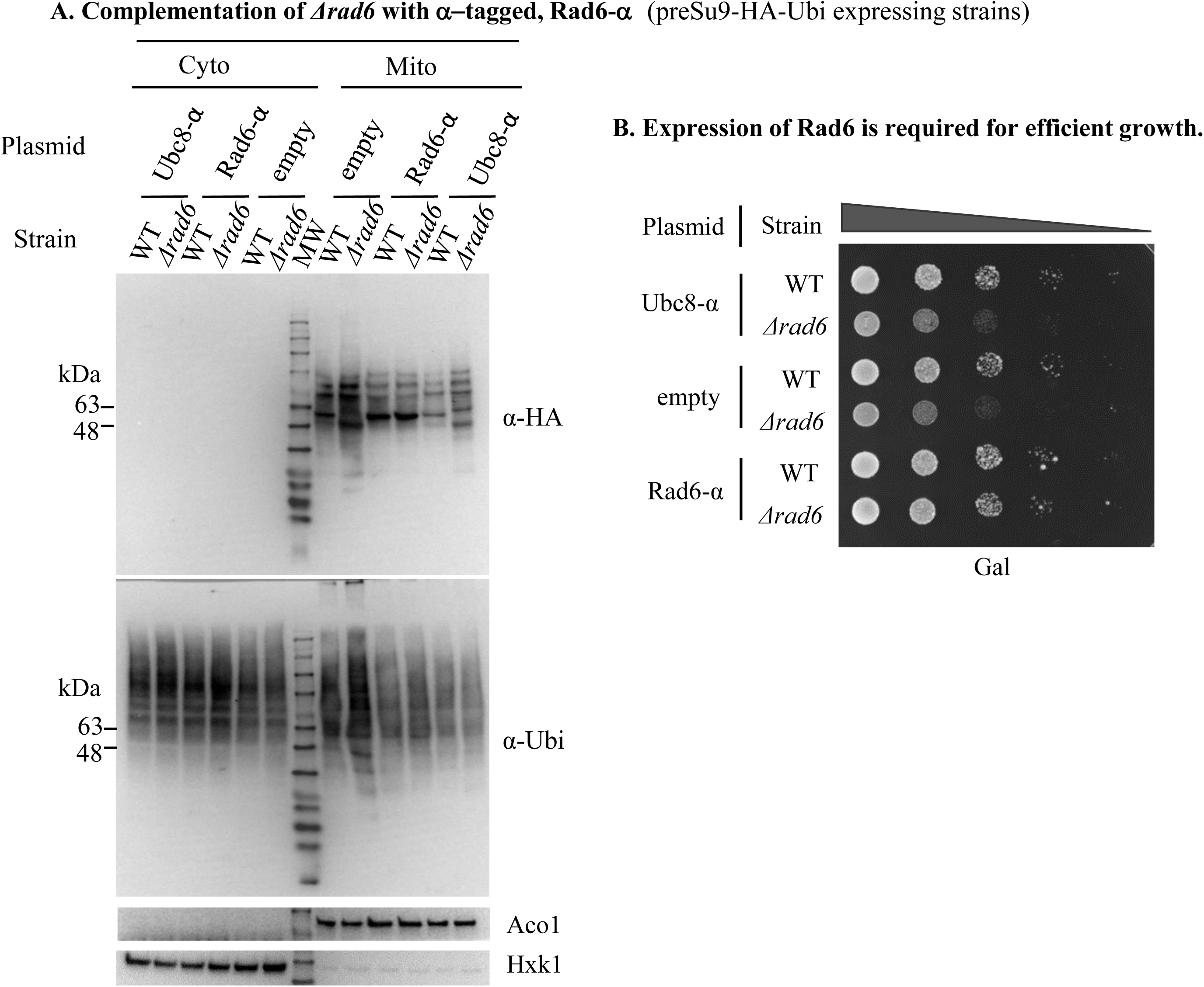
RAD6 deficiency can be complemented by an active Rad6 expressed from a plasmid. **(A) Rad6-α restores the preSu9-HA-Ubi ubiquitination pattern of the RAD6 KO cells.** Rad6-α, Ubc8-α, or empty plasmid constructs were introduced into preSu9-HA-Ubi expressing WT or *Δrad6* cells. Subcellular fractions were analyzed by WB as in Figure 5. **(B) Rad6-α restores efficient growth in *Δrad6* cells.** The growth of the strains analysed in Figure (A) were examined by a growth drop assay.

The altered pattern, or the missing major band signal of preSu9-HA-Ubi conjugates in RAD6 KO cells, suggests that Rad6 is active in mitochondria. To verify this assumption, we examined if Rad6 which is specifically targeted to mitochondria, can restore the normal preSu9-HA-Ubi conjugate pattern of mitochondria in *Δrad6* cells. Initially, a preSu9-Rad6-α expression vector was constructed. However, α- complementation shows incomplete import of preSu9-Rad6-α into mitochondria, as a light blue colour was observed with the ωc strain of both WT and *Δrad6* background (Figure 7A). Therefore, Rad6-α fused to the preSu9 MTS (at its N terminus) was also fused to a degron, SL17, at its C terminus (preSu9-Rad6-α-SL17). The SL17 degron element makes the fusion protein in the cytosol a ubiquitin-proteasome substrate [31] and therefore is destined for direct degradation of the preSu9-Rad6-α-SL17 in the cytosol, whereas the same protein imported into mitochondria is protected. The α- complementation assay demonstrates that preSu9-Rad6-α-SL17 was only detected in mitochondria, indicating sole distribution of preSu9-Rad6-α-SL17 to mitochondria (Figure 7A).

**Figure 7:**
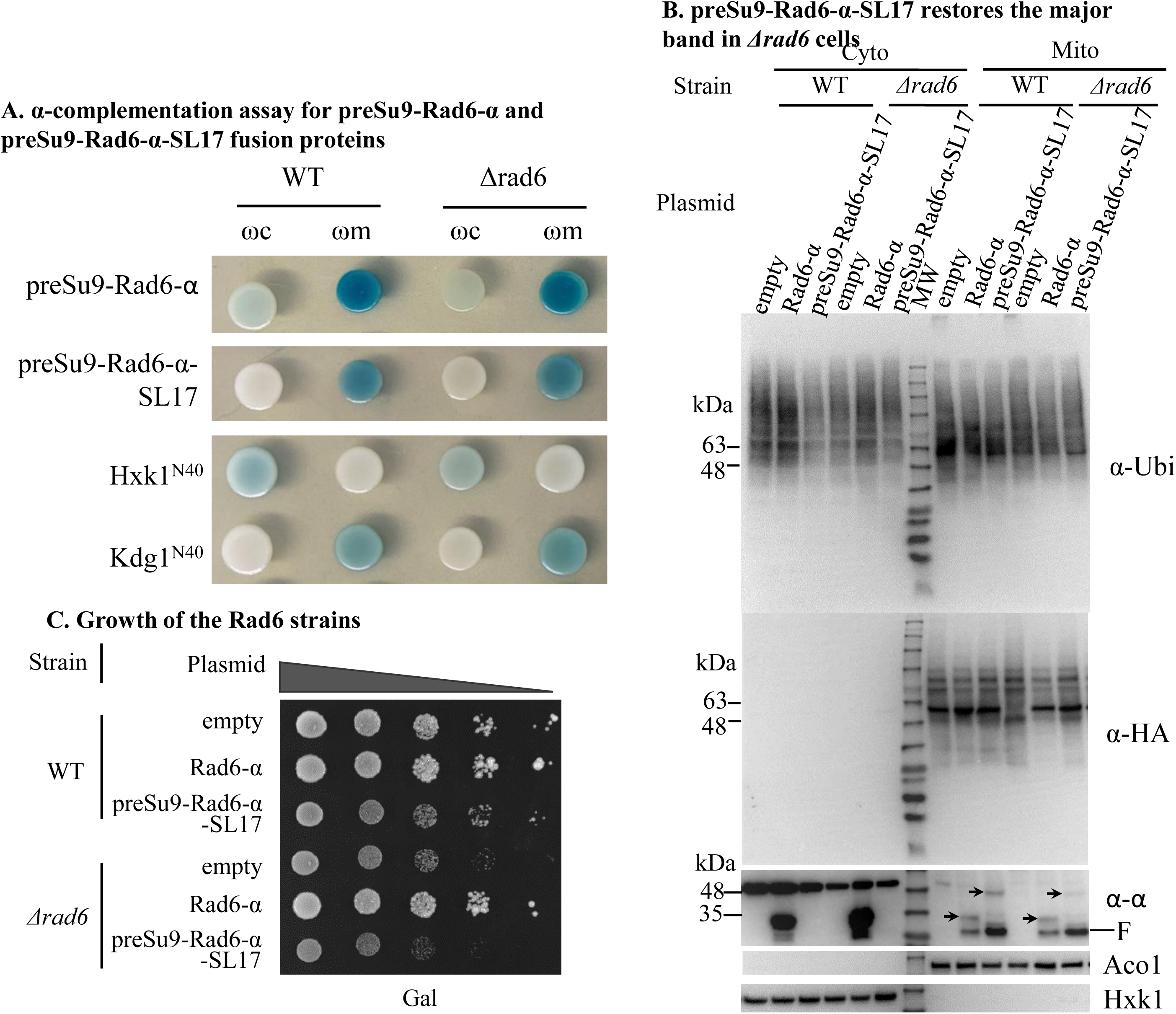
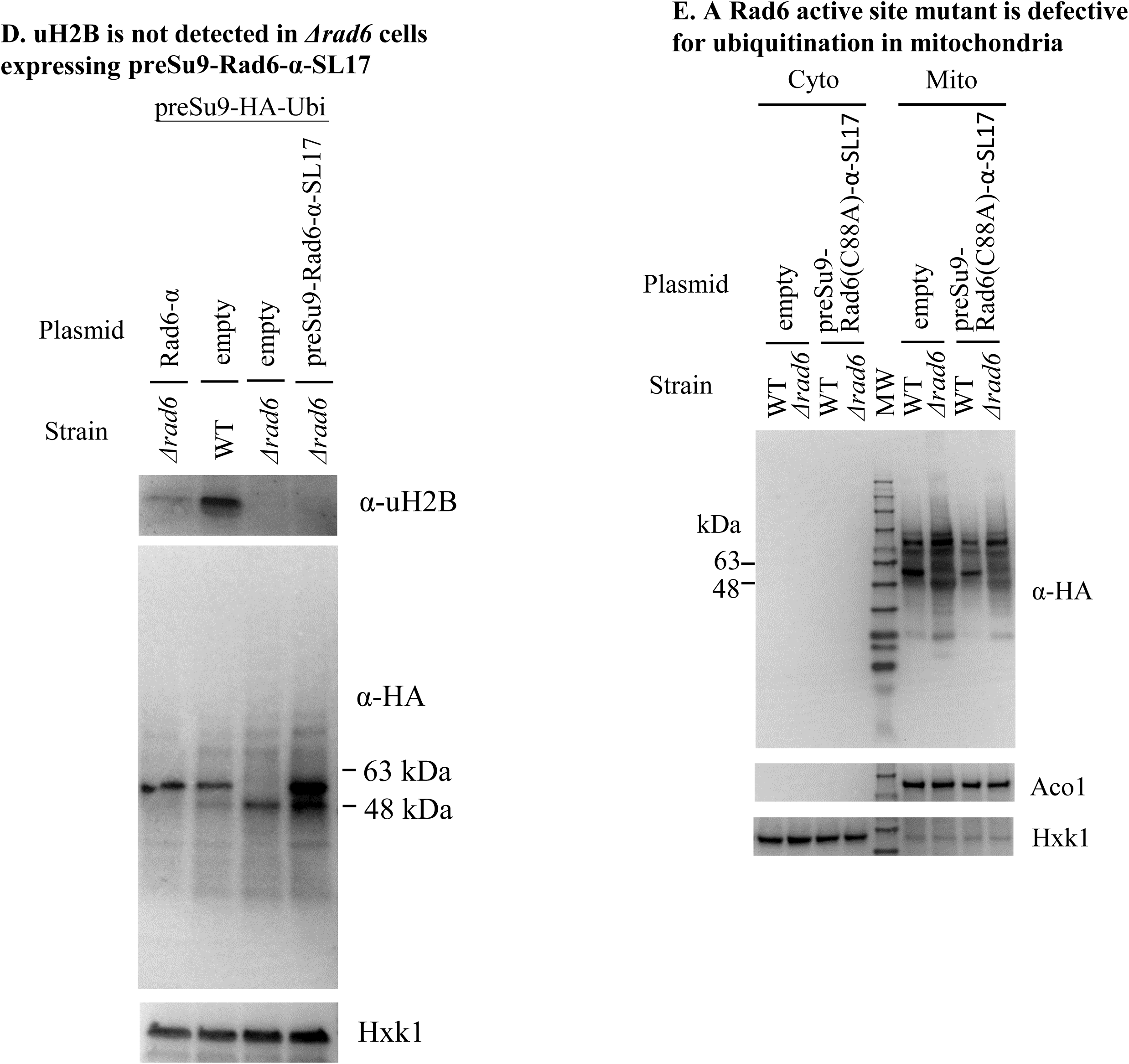
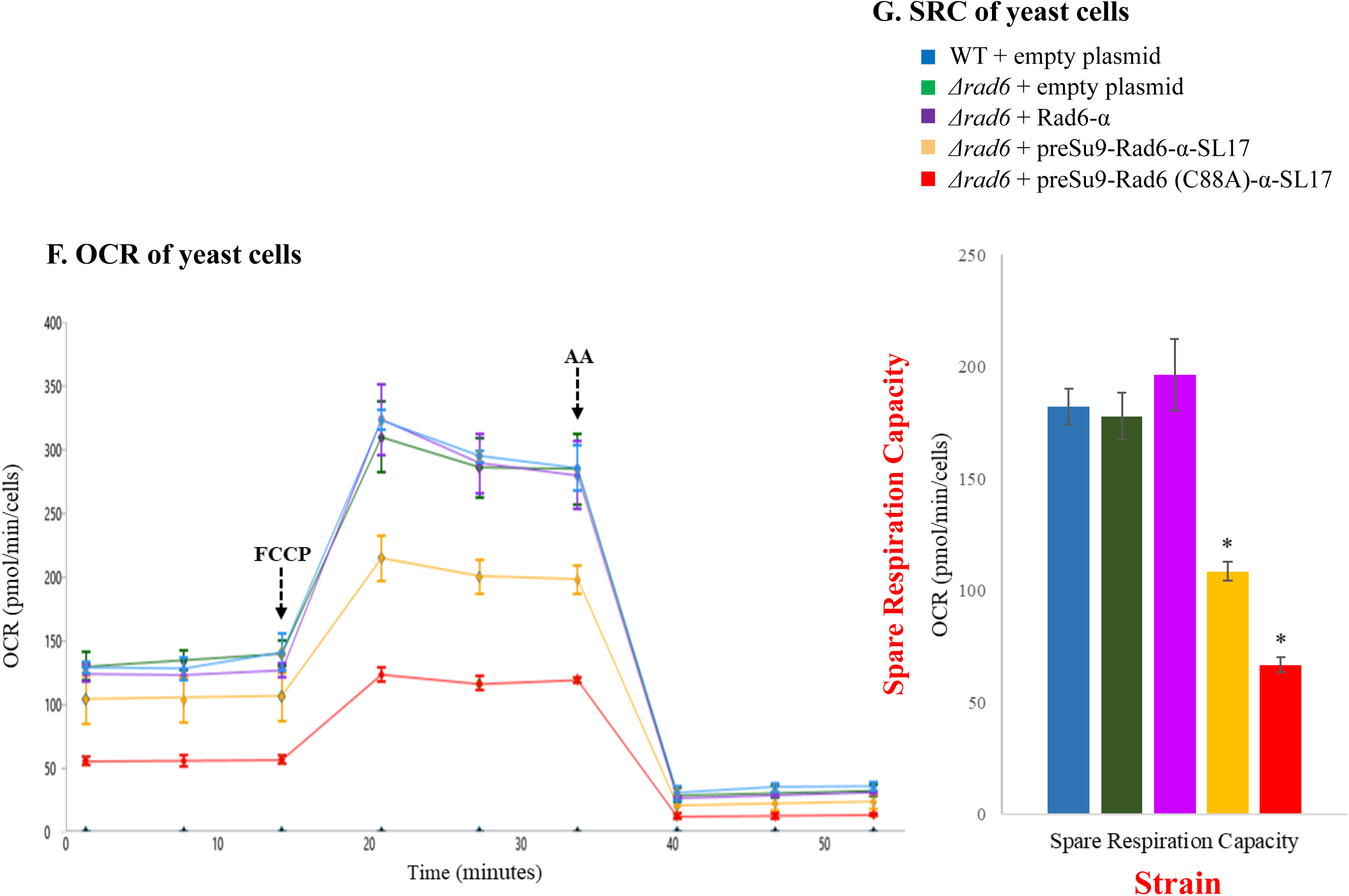
Rad6 is active in protein ubiquitination in mitochondria. **(A) α-complementation assay for preSu9-Rad6-α and preSu9-Rad6-α-SL17 fusion proteins.** N terminal 40 amino acids of cytosolic protein Hxk1 (Hxk1^N40^) and mitochondrial protein Kgd1 (Kgd1^N40^) fused to the α- fragment are cytosolic and mitochondrial controls. **(B) preSu9-Rad6-α-SL17 restores the major band in *Δrad6* cells.** preSu9-HA-Ubi expressing cells harboring an empty plasmid, a plasmid encoding Rad6-α or preSu9-Rad6-α-SL17 were subjected to subcellular fractionation followed by WB of the indicated subcellular fractions. Cytosolic and mitochondrial fractions (Cyto and Mito) were probed with α-HA, α-Ubi and α-alpha (α-α). Aco1, mitochondrial marker; Hxk1, cytosolic marker. Line arrows indicate the band of Rad6-α or preSu9-Rad6-α-SL17. F, fragment of Rad6 fusion proteins. **(C) Growth of the Rad6 related strains.** Strains analyzed in Figure 7B were examined for growth by a drop assay on semi-solid medium. **(D) uH2B was not detected in preSu9-Rad6-α-SL17 expressing *Δrad6* cells.** Whole cell extracts were prepared from strains harboring the indicated plasmids and analyzed by WB for the level of monoubiquitinated H2B (uH2B), HA and Hxk1 (loading control). **(E)** Appearance of preSu9-HA-Ubi conjugates major band requires the E2 enzymatic activity of Rad6. Rad6 point mutation construct, preSu9-Rad6 (C88A)-α-SL17, or empty plasmid was transformed into preSu9- HA-Ubi harbouring WT or *Δrad6* cells. Cytosolic and mitochondrial subcellular fractions of the indicated transformants were analysed for the conjugate patterns by WB using α-HA antibody. **(F and G) Oxygen Consumption Rate (OCR) and Spare Respiration Capacitary (SPC) of yeast cells.** Yeast strains were grown to mid-log phase in galactose media and shifted to assay media with 2% ethanol-acetate. The oxygen consumption rate of each yeast strain was measured using the Seahorse XFe96 Analyzer. The cell density was 5×10^5^cells per well. The OCR values are calculated per OD_600_. The Spare Respiration Capacitary (SPC) was calculated from the difference between the basal respiration and maximal respiration capacity (following addition of FCCP). Injections of FCCP and antimycin A (AA) are indicated with arrows. All data represent the mean ± SD of biological triplicates *: p<0.05. The yeast strains analysed were WT + empty plasmid (), *Δrad6* + empty plasmid (), *Δrad6* + Rad6-α (), *Δrad6* + preSu9-Rad6-α-SL17 (), *Δrad6* + preSu9-Rad6 (C88A)-α-SL17 ().

The presence of preSu9-HA-Ubi conjugates in mitochondrial fractions was tested by WB of extracts of *Δrad6* cells expressing preSu9-Rad6-α-SL17. Figure 7B (second panel) shows that the major band missing in the *Δrad6* cells was restored by preSu9- Rad6-α-SL17; moreover, the strength of the restored major band seemed to be even stronger than that of WT cells harbouring an empty plasmid or *Δrad6* cells harbouring Rad6-α. This suggests that preSu9 MTS, in the preSu9-Rad6-α-SL17, has an effect of enhanced targeting of Rad6 to mitochondria. Using antiserum against the alfa-tag, only preSu9-Rad6-α-SL17 in the mitochondrial fraction was detected (Figure 7B, 3^rd^ panel), consistent with the α-complementation assay in Figure 7A. Rad6-α was detected as a mitochondrially eclipsed protein (Figure 7B, 3^rd^ panel), like the endogenous Rad6, which in fact is expressed at very low levels and can hardly be detected in the mitochondrial fraction by WB (Figure S5). Importantly, unlike Rad6-α, preSu9-Rad6-α-SL17 cannot restore the full cell growth of *Δrad6* cells on galactose as the carbon and energy source, indicating sole distribution of preSu9-Rad6-α-SL17 to mitochondria and that Rad6 outside mitochondria is required to maintain normal cell growth under specific conditions (Figure 7C). Taken together these data suggest that Rad6 is eclipsed distributed in mitochondria and is active in the organelle.

Rad6 is known to direct mono-ubiquitination of histone H2B, which is involved in regulating gene transcription in the nucleus [32]. Mono-ubiquitinated H2B (uH2B) was detected by WB in both WT cells and *Δrad6* cells expressing Rad6-α but not in *Δrad6* cells (Figure 7D, top panel, compare the three left lanes). Importantly, uH2B was not detected in preSu9-Rad6-α-SL17 expressing *Δrad6* cells (Figure 7D, top panel, right lane), while the major band strength was again detected to be stronger than the native level (Figure 7D, middle panel). This indicates that the transcription of the major band protein(s), is not dependent on Rad6 mediated mono-ubiquitination of histone H2B and further supports the active role of Rad6 in mitochondria.

### Rad6 UBC activity is required for protein ubiquitination in mitochondria

To confirm that it is the enzymatic activity of Rad6 in mitochondria that is required for the formation of the preSu9-HA-Ubi conjugates major band, we constructed the expression vector preSu9-Rad6(C88A)-α-SL17 with a point mutation, C88A, in the Rad6 ubiquitin-conjugating (Ubc) active site. Rad6 with this point mutation is defective in Ubc activity [33]. Figure S5 (left panel) and Figure S6 show the expression and targeting of preSu9-Rad6(C88A)-α-SL17 to mitochondria. However, preSu9- Rad6(C88A)-α-SL17 fails to restore the preSu9-HA-Ubi conjugates major band in *Δrad6* cells (Figure 7E), proving the requirement for Ubc activity of Rad6 in mitochondria. In other words, Rad6 activity is required for protein ubiquitination in mitochondria.

Interestingly, in respirometry experiments to measure the OCR (Oxygen Consumption Rate), we found that *Δrad6* expressing preSu9-Rad6α-SL17 exhibited a ∼40% drop in OCR and SRC (Spare Respiration Capacity, Fig. 7F and 7G, compare the WT and *Δrad6* + preSu9-Rad6-α-SL17). Furthermore, *Δrad6* expressing preSu9-Rad6(C88A)- α-SL17 exhibited an even stronger ∼70% drop in OCR. We therefore find that targeting overwhelmingly high protein levels of Rad6 to mitochondria can affect OCR regardless of whether it is active or not. This may be explained by protein-protein interactions that Rad6 undergoes in mitochondria.

We subjected the major band to MS analysis with the assumption that protein(s) representing the major band are hopeful candidate substrate(s) of Rad6. Specifically, after proteins of mitochondrial fractions of preSu9-HA-Ubi expressing WT and *Δrad6* cells were separated on SDS-PAGE, the region (between 48 and 63 kDa corresponding to the migration position of the major band) was excised for the two separated samples and processed for MS. Table S2 summarizes the ubiquitinated proteins detected in each sample by MS. One of the proteins that was detected was Ilv5, which appears to be ubiquitinated at different sites in the WT versus the *Δrad6* cells and at a higher frequency in the WT cells. This candidate together with other proteins (against which antibodies are available in our laboratory), such as Aco1, Hsp60, Fum1, and Idh1, were subjected to analysis. Mitochondrial lysates of cells expressing preSu9-HA-Ubi were subjected to IP using anti-HA magnetic beads and proteins eluted from the beads were analysed by WB with the antibodies of specific proteins referred to above. We found no significant differences between WT and *Δrad6* cells in terms of the amount of modified Ilv5 which could of explain changes in the major band. Nevertheless, the detection of the modified form of matrix proteins, Ilv5 and Aco1, in the preSu9-HA-Ubi IPed samples coincides with the notion of ubiquitination in mitochondria (Figure S7).

### The Rad6 N terminal is required for its targeting to and activity in mitochondria

Recently we performed a screen for novel dual targeted *S. cerevisiae* mitochondrial proteins by using the α-complementation assay (see above, manuscript in preparation). The N-terminal 11 amino acids of Rad6 are predicted to be critical residues for the high MitoProt II score, 0.9876. The MitoProt II score for Rad6 without the N-terminal 11 amino acids dramatically drops to around 0.1 (Figure 8A). Besides being suspected to have a mitochondrial targeting function, the N-terminal of Rad6 may also be essential for its E2 activity. In this regard, the N-terminal 15 residues of Rad6, are almost identical between species [34], and it has been reported to be required for its functions in different pathways such as sporulation, N-end rule protein degradation, and DNA repair [35]. To verify if the N-terminal of Rad6 is required for its targeting to and enzymatic activity in mitochondria, Rad6-α truncated at its N-terminus by 11 amino acids (Rad6^ΔN11^-α) was constructed. This protein is not targeted to mitochondria according to the α-complementation assay (Figure 8B, upper panel) and it lacks the ability to restore the ubiquitination pattern in mitochondria (Figure 8C).

**Figure 8:**
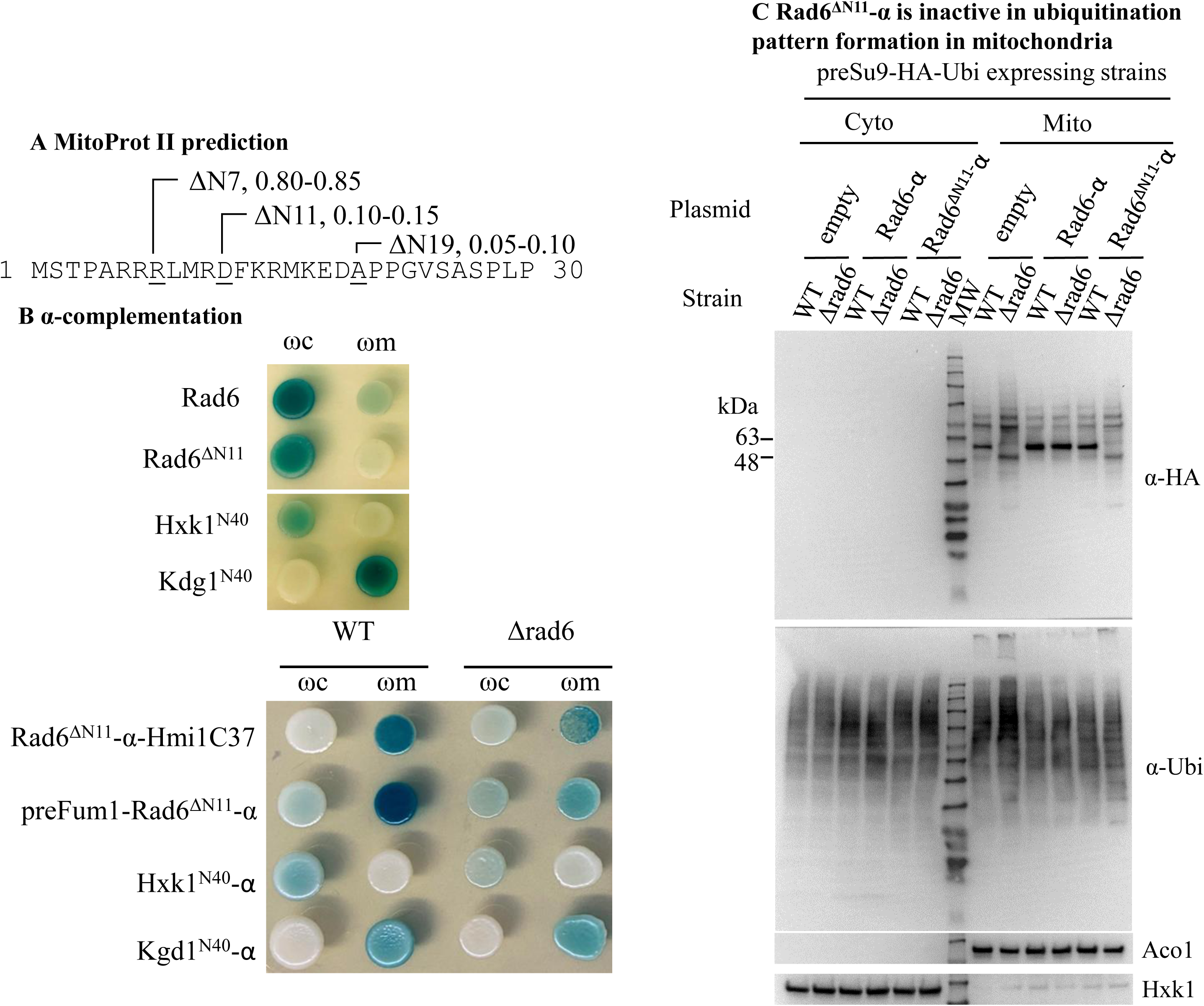
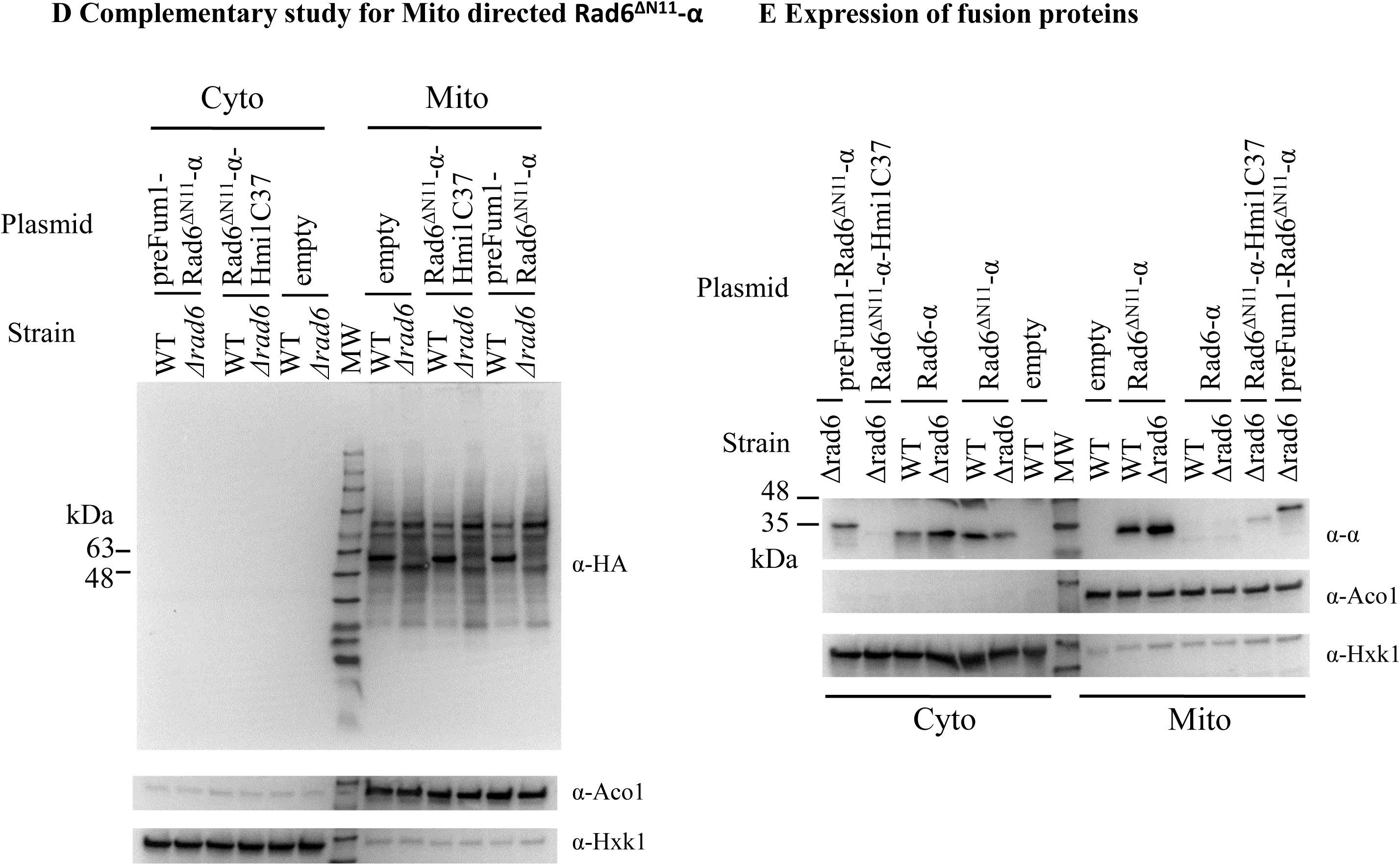
N-terminal amino acids of Rad6 is necessary for its activity in mitochondria. **(A) MitoProt II analysis of Rad6 N-terminal truncations at different positions.** The Mitoprot II scores of Rad6 with its N-terminal truncated at 7, 11, or 19 amino acids (ΔN7, ΔN11, or ΔN19) and a methionine at the N terminal. **(B) α-complementation of Rad6 N-terminally truncated by 11 amino acids (Rad6^ΔN11^-α, and MTS directed Rad6^ΔN11^-α).** Rad6^ΔN11^-α N-terminally fused to the MTS of Fum1 or C-terminally to the MTS of Hmi1 (the last 37 amino acids of the protein) is designated as preFum1-Rad6^ΔN11^-α and Rad6^ΔN11^-α-Hmi1C37, respectively. **(C) Rad6^ΔN11^-α can not restore the normal ubiquitination pattern in mitochondria in *Δrad6* cells.** preSu9-HA-Ubi conjugates signal was investigated for cytosolic and mitochondrial fractions of WT or *Δrad6* cells harbouring indicated plasmids by WB. Blot was also probed for control proteins, Ubiquitin (Ubi), Aco1 (mitochondrial marker) and Hxk1 (cytosolic marker). **(D) MTS directed Rad6^ΔN11^ can not restore the normal ubiquitination pattern in mitochondria of *Δrad6* cells.** preSu9-HA-Ubi conjugates signals were investigated for cytosolic (Cyto) and mitochondrial (Mito) fractions of WT and *Δrad6* cells harbouring MTS directed Rad6^ΔN11^-α or an empty plasmid by WB with α-HA antibodies. Aco1, mitochondrial marker; Hxk1, cytosolic marker. **(E) Distribution of MTS directed Rad6^ΔN11^-α between the cytosolic and the mitochondrial fraction.** Fusion proteins were probed using antibody against α-tag, α- α. Aco1, mitochondrial marker; Hxk1, cytosolic marker.

In addition, we constructed Rad6^ΔN11^-α which is N-terminally fused to the MTS of Fum1 or C-terminally to the matrix targeting signal of Hmi1 (the last 37 amino acids of the protein [36]). While the Rad6^ΔN11^-α is not targeted to mitochondria, the two matrix-directed Rad6^ΔN11^-α fusion proteins (respectively designated as preFum1-Rad6ΔN11- α and Rad6^ΔN11^-α-Hmi1C37) are targeted to mitochondria matrix according to α- complementation assay (Figure 8B, bottom panel). Regarding the Rad6 activity, the WB shows that neither preFum1-Rad6^ΔN11^-α nor Rad6^ΔN11^-α-Hmi1C37, can restore the preSu9-HA-Ubi conjugates major band in *Δrad6* cells (Figure 8C), suggesting that the N-terminal 11 amino acids of Rad6 are indispensable for the mitochondria relevant activity. These results clearly indicate that the Rad6 amino terminus is involved in both targeting to and activity of the protein in the mitochondrial matrix.

## DISCUSSION

Mitochondria evolved from an α-proteobacterium-like prokaryotic cell that was engulfed by an Asgard Archaea-like ancestor of eukaryotic cells [37]. Ubiquitination is a type of post translational modification unique to eukaryotic cells even though some ubiquitination system components can be detected in certain Archaea [38]. Since mitochondria contribute a small fraction to the total cellular protein mass (less than 1/15 as found in our study), ubiquitination in mitochondria can be easily overlooked. Following an initial suggestion for ubiquitination in human mitochondria [1], we provide systematic proof for ubiquitination in *S. cerevisiae* mitochondria:

(1) The α-complementation screen detected ubiquitination system components in the matrix of *S. cerevisiae* mitochondria (E1, E2, E3, etc).
(2) Conjugates of MTS directed HA-Ubi (preSu9-HA-Ubi) are detected only in mitochondrial fractions.
(3) Conjugate signals of preSu9-HA-Ubi in isolated mitochondria are resistant to external trypsin treatment.
(4) preSu9-HA-Ubi conjugate levels in mitochondria are not elevated by proteasome inhibition (MG132 treatment).
(5) Immunoprecipitation (IP) of preSu9-HA-Ubi conjugates is highly enriched for mitochondrial matrix proteins.
(6) The E3 RAD6 absence from mitochondria causes an altered ubiquitination pattern of proteins in the organelle.
(7) Rad6 inactivated by the point mutation C88A (active site) does not complement the effect the Rad6 KO on the pattern of ubiquitinated proteins in mitochondria.

Where does ubiquitination of mitochondrial proteins occur: the matrix, membranes or intermembrane space? In our study, MTSs, which are matrix targeting signals (preSu9 and preFum1), are used to direct import of HA-Ubi into the mitochondrial matrix [39, 40]. In particular preSu9 (1-69) is established as a highly efficient matrix-targeting signal in yeast [20, 41, 42]. Therefore, we reason that MTS-HA-Ubi conjugates resistant to external trypsin treatment of isolated mitochondria are formed in the matrix. Furthermore, the preSu9-Rad6-α-SL17 construct, which is solely targeted to mitochondrial matrix, can restore the major band of preSu9-HA-Ubi conjugates in mitochondria in *Δrad6* cells. These observations taken together indicate ubiquitination in the matrix of *S. cerevisiae* mitochondria.

We have several components of the ubiquitination system, including E1, E2, E3, etc., testing positive for mitochondrial targeting in our α-complementation assay. Why was the mitochondrial targeting of components of ubiquitination system overlooked in previous studies? The principal reason lies in eclipsed mitochondrial targeting. In other words, a predominant portion of the dual targeted ubiquitination system proteins is localized outside mitochondria with only minute amounts being targeted to the organelle, thereby obscuring detection of the latter. And in fact, the E2 Rad6, which is specifically investigated in this study, is shown to be eclipsed distributed in mitochondria. In addition, in our proteome-scale analysis of proteins relative abundance between mitochondrial and cytosolic fractions (unpublished data), 12 out of the 15 ubiquitination system proteins testing positive in our α-complementation (Figure 1B and C) are detected, and 11 are indicated with eclipsed amounts in mitochondrial fractions, including Uba1, Stp22, Cdc34, Ubc8, Rad6, Mms2, Ubc13, Cdc16, Mpe1, Ubp13, and Duf1. One could argue that such small amounts of proteins inside mitochondria are some kind of “contamination” with no real function or consequence. Here we show that absence of active Rad6 in mitochondria affects the pattern of ubiquitination in the organelle, proving the function of this eclipsed protein in mitochondria.

It is worth pointing out that proteasome subunits are not detected in mitochondria. In the present study the proteasome activity is not detected on ubiquitinated proteins in the organelle: MG132 does not change the pattern of ubiquitinated mitochondrial proteins. Furthermore, the proteome associated degron SL17 does not lead to the clearance of preSu9-Rad6-α-SL17 from mitochondria. Accordingly, one of the interesting directions for future studies will be to ask whether mitochondrial proteases, such as the matrix facing Yta10–Yta12 m-AAA protease on the inner membrane and the matrix protease Pim1 [43], assume the quality control function of degrading certain ubiquitinated mitochondrial proteins.

This study reveals a novel role for the activity of Rad6, which is eclipsed in mitochondria, affects the pattern of ubiquitinated proteins (specifically conjugated by preSu9-HA-Ubi) in the organelle. It is important to stress that the active site mutant of Rad6, Rad6(C88A), cannot complement the Rad6 effect on the pattern of ubiquitinated proteins in mitochondria. Ilv5, identified as a ubiquitination substrate by mass spectrometry of the extracted major band by us, was however disappointing since the level of the modified protein in mitochondria appeared similar in the WT and *Δrad6* cells. Nevertheless, preSu9-HA-Ubi conjugates signals in Rad6 deficient mitochondria suggests that Rad6 is not the only ubiquitination cascade protein active in the organelle. In addition, our α-complementation screen suggests import of other ubiquitination cascade proteins into mitochondria.

There are numerous possibilities in terms of targeting components of the ubiquitination system to mitochondria. Nevertheless, regarding Rad6, both the program, MitoProt II, prediction and the experimental validation suggest the MTS function of the N terminal of Rad6. The Rad6 protein is predicted to contain an N-terminal MTS like sequence (MitoProt II); deletion of the first 11 amino acids of Rad6, which covers a critical region of the predicted MTS, is enough to block its import into the mitochondrial matrix. In addition, this truncated version of Rad6 directed to mitochondria lacks the activity to restore the WT ubiquitination pattern in mitochondria of *Δrad6* cells. The observed requirement of Rad6 N terminal for its activity in, and targeting to, mitochondria may explain the high conservation of the N-terminal 15 residues of Rad6 between species [34].

A question for which we have no answer is, with which E3(s) does Rad6 interact while functioning in mitochondria? Rad6 is an E2 and is expected to interact with E3s. In fact, in the present study, we tested the deletion strains of five E3 (related) proteins that are known to interact with Rad6 as part of its cytosolic/nuclear function, as well as an additional 13 E3s, but none was detected to influence the pattern of protein ubiquitination in mitochondria. While there are about 80 E3s awaiting to be fully tested, it is also possible that ubiquitination of the Rad6 substrate(s) in mitochondria is not E3 dependent, as yeast Rad6 can modify substrates on its own [44, 45].

This study of ubiquitination in *S. cerevisiae* mitochondria reveals a new perspective regarding mitochondrial and ubiquitination research. We believe that this study will have a profound impact on revealing how ubiquitination of proteins in mitochondria regulates mitochondrial/cellular functionality. It is important to study ubiquitination in mitochondria of other eukaryotic organisms. Such studies may allow us to gain insights into how during evolution, the endosymbiotic organelle, mitochondria, acquired the ubiquitination system and how this was integrated into the needs of the whole cell.

## METHODS

### Growth conditions of yeast cells

Cells were grown at 30 °C in rich medium containing 1% (w/v) yeast extract, 2% (w/v) peptone, and 2% (w/v) glucose or galactose, or in synthetic depleted (SD) minimal medium containing 0.67% (w/v) yeast nitrogen base and 2% (w/v) galactose or glucose, supplemented with appropriate amino acids (all the components of complete amino acid mixture except the ones corresponding the auxotrophic markers of the plasmids contained in yeast cells). Unless specified otherwise, cells were cultured in medium with galactose as the carbon and energy source. For expression of preSu9- HA-Ubi driven under *MET25* promoter, amino acid methionine should be absent in the medium. 15% agar was added for agar plates.

### α-Complementation assay

Yeast cells co-expressing α-fusion proteins in combination with ωc or ωm were suspended in distilled H_2_O. Suspended cells were then dropped onto 90 mg/L X-Gal plates. Results were collected after incubation at 30°C for ∼7 days.

### Subcellular fractionation

Induced yeast cultures were grown to OD_600_ of around 1.5. Subcellular fractionations for the cells collected were performed as previously described with some modifications [46]. Spheroplasts were prepared in the presence of Zymolyase 20T (MP Biomedicals, Irvine, CA). 2-3 ml of homogenization buffer (0.6 M sorbitol, 20 mM HEPES, pH7.5) supplemented with 1x Protease Inhibitor Cocktail (P8215, Sigma) and 0.5 mM PMSF was applied to spheroplasts prepared from 100 OD_600_ units of yeast cells for homogenization. The homogenized spheroplasts cleared of cell debris by subsequent 1500x g centrifugation for 6 min are defined as total fractions. 10000x g centrifugation for 10 min was used to sperate mitochondria from the total fractions. The post mitochondrial supernatant and the isolated mitochondria, which were resuspended after washing are defined as cytosolic and mitochondrial fractions, respectively. Subcellular fractionations were assayed for cross contaminations using antibodies against mitochondrial marker, α-Hsp60, α-Tom70 and/or α-Aco1, and cytosolic marker, α-Hxk1.

### Yeast oxygen consumption rate measurements using Seahorse XFe96 analyzer

Cells were grown in galactose media at 30°C with agitation to 1.0 OD at 600nm. Next, cells were washed, counted and resuspended in the assay medium (0.67% yeast nitrogen base, 2% potassium acetate and 2% ethanol). The cells were then diluted and 180 µl of resuspended cells seeded in pre-coated Poly-L Lysine (22 µl of 0.1 mg/ml per well) XFe96-well microplates (Agilent Seahorse Technologies) at 5×10^5^cells per well. After seeding, cells were centrifuged at 500g for 3 min and incubated at 30°C for 30 min in the assay medium prior to measurements. The Agilent Seahorse XFe96 sensor cartridge was consecutively hydrated at 30°C with sterile water for overnight and in XF Calibrant for 1 hr following the manufacturer’s instructions. After that, compound solutions of 20 µM FCCP and 2.5 µM Antimycin-A were loaded into the corresponding ports, A and B using the well labels (lettered A-H) on the Seahorse XF assay cartridge. The Seahorse XFe96 Analyzer was set to maintain the temperature at 30°C. Both the mixing time and measuring time were set to 3 min in each cycle. The oxygen consumption rate is shown as picomole oxygen per minute per cell (pmol/min/cell). Data represent the average of three independent experiments for each strain.

### Immunoprecipitation

Mitochondrial pellet obtained through subcellular fractionation, was lysed in Pierce^TM^ IP Lysis Buffer. Pierce™ Anti-HA Magnetic Beads was used to immunoprecipitate proteins from mitochondrial lysates. IP were manually performed according to the anti-HA beads product instructions with some modifications indicated elsewhere.

### Mitochondrial trypsinization

Mitochondria pellets obtained from approximately 55 OD_600_ units of yeast cells through subcellular fractionation were suspended in 150 μl of homogenization buffer and incubated for 30 min on ice with 2 μl of distilled H_2_O or 1 μg/μl trypsin. Trypsinization was stopped by addition of protein loading dye and subsequent boiling at 100 °C for 11 min.

### MG132 induced proteasome inhibition of living yeast cells

125 OD_600_ units of yeast cells (*Δpdr5* background) grown to logarithmic phase in galactose minimal medium were collected and then suspended in 30 ml galactose rich medium (YPGal) with or without 60 μM of MG132. Cells continued to grow in YPGal until the OD was doubled, before being collected for subcellular fractionation.

### Mass spectrometry (MS)

After proteins in samples were trypsinated, the tryptic peptides were subjected to liquid chromatography–mass spectrometry (LC–MS) analysis. Sample trypsinization and MS were processed at Protein and Proteomics Centre (PPC), NUS Biological Sciences.

---

**Figure S1:**
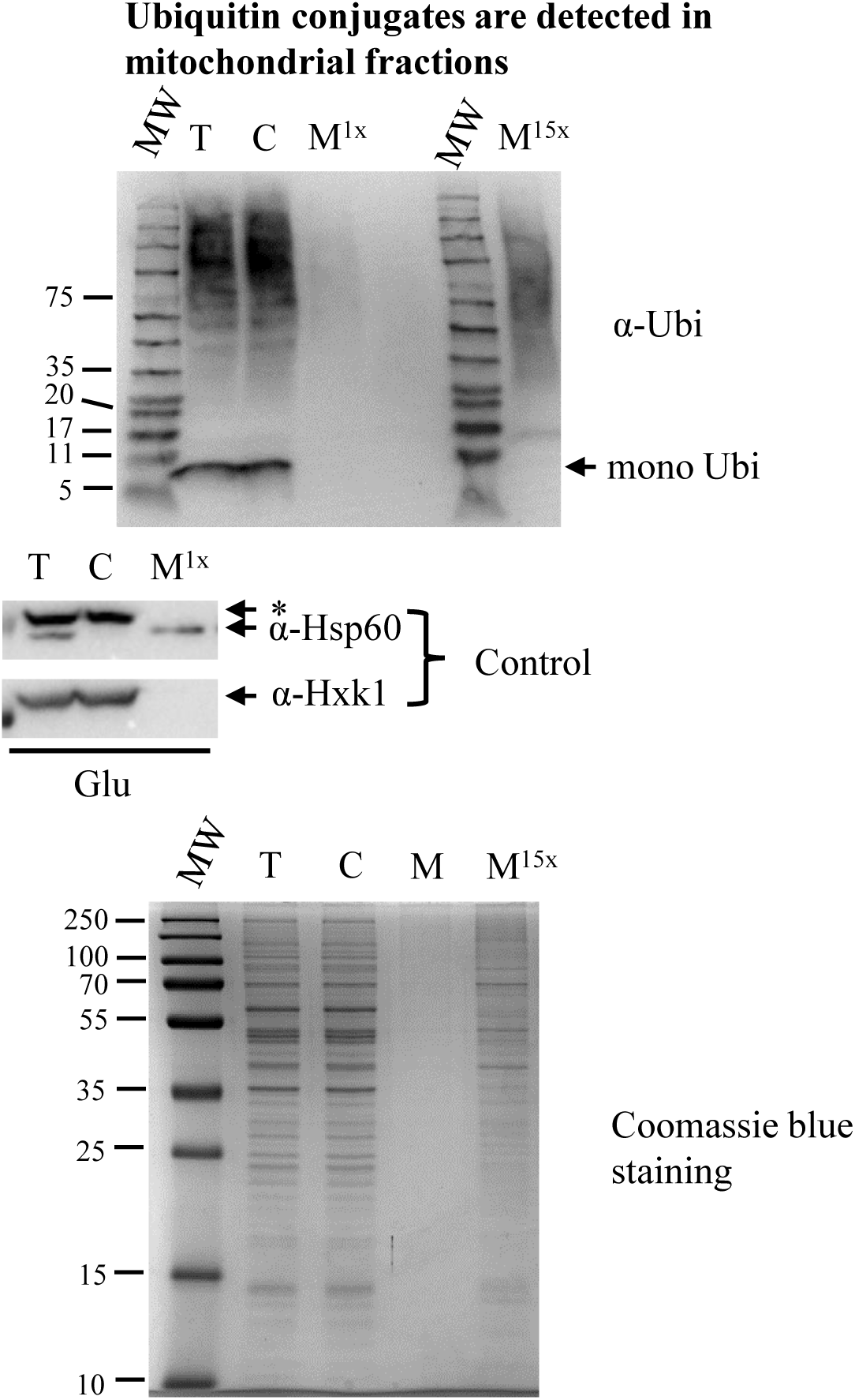
Ubiquitin conjugates are detected in mitochondrial fractions. **Top Panel: Subcellular fractions probed for ubiquitin.** Subcellular fractions prepared from cells grown to logarithmic phase in glucose minimal medium were subjected to analysis as described in the legend of Figure 2. **Middle Panels:** T, C, and M fractions were probed with control antibodies against Hsp60 and Hxk1, which are mitochondrial and cytosolic markers, respectively. *, The upper bands detected in T and C fractions are non-specific. **Bottom Panel:** Subcellular fractions were subjected to 12.5% SDS-PAGE and Coomassie Blue staining.

**Supplementary Figure S2:**
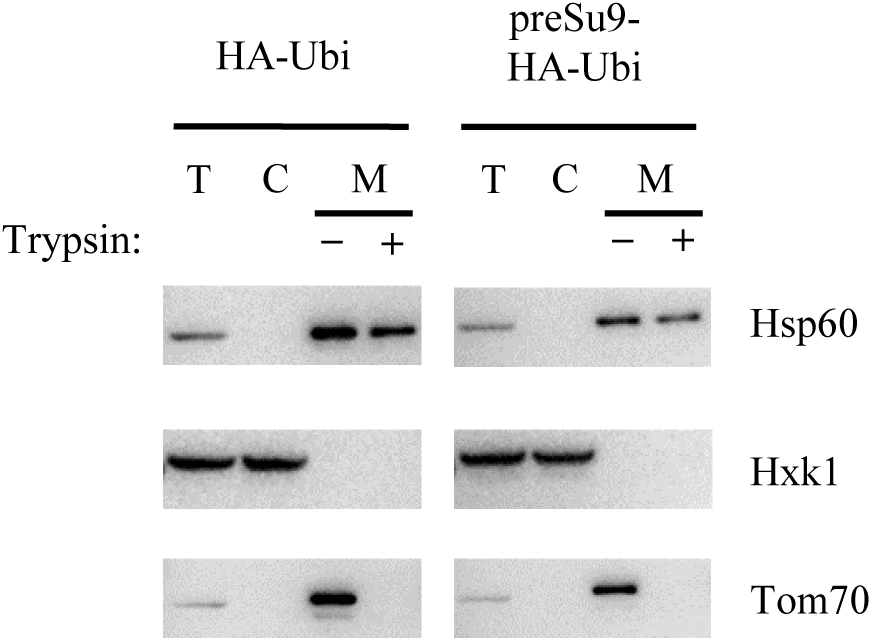
Fractionation controls for. **Figure 3D** Quality of the fractionation was monitored by mitochondrial marker - Hsp60 and cytosolic marker - Hxk1. T, total fraction; C, cytosolic fraction; M, mitochondrial fraction. Tom70, a mitochondrial outer membrane protein as a marker exposed to trypsin treatment.

**Figure S3:**
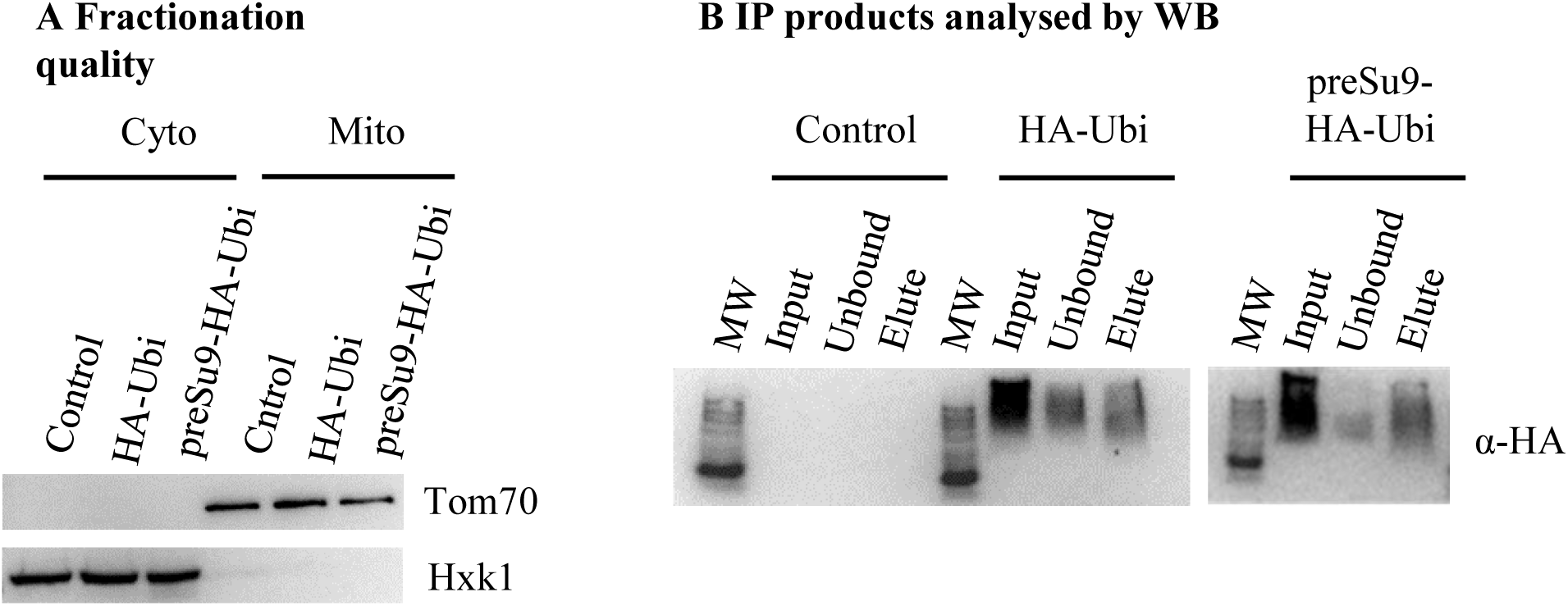
Immunoprecipitation (IP) for ubiquitinated conjugates marked by HA. **(A) Quality of mitochondrial isolate.** Mitochondrial (Mito) and cytosolic (Cyto) fractions corresponding to approximately 2x and 1x cells respectively were loaded for WB to verify the efficiency in separation between the indicated two types of fractions. Hxk1, cytosolic marker; Tom70, mitochondrial marker. **(B) IP products analysed by WB.** Three IPs (anti-HA) were conducted in parallel using mitochondrial lysates described in (A). 300 μl of lysate corresponding to ∼180 OD_600_ units of yeast cells were applied for each IP. 0.1 M glycine solution, pH 2.0, was used to elute proteins from immunoprecipitated beads. Lysates before (Input) and after (Unbound) incubation with anti-HA magnetic beads were loaded in the same volume for WB. One 11^th^ of each elution was loaded for WB. Precipitation of ubiquitinated conjugates marked by HA was confirmed using primary antibody against HA (α-HA).

**Supplementary Figure S4.**
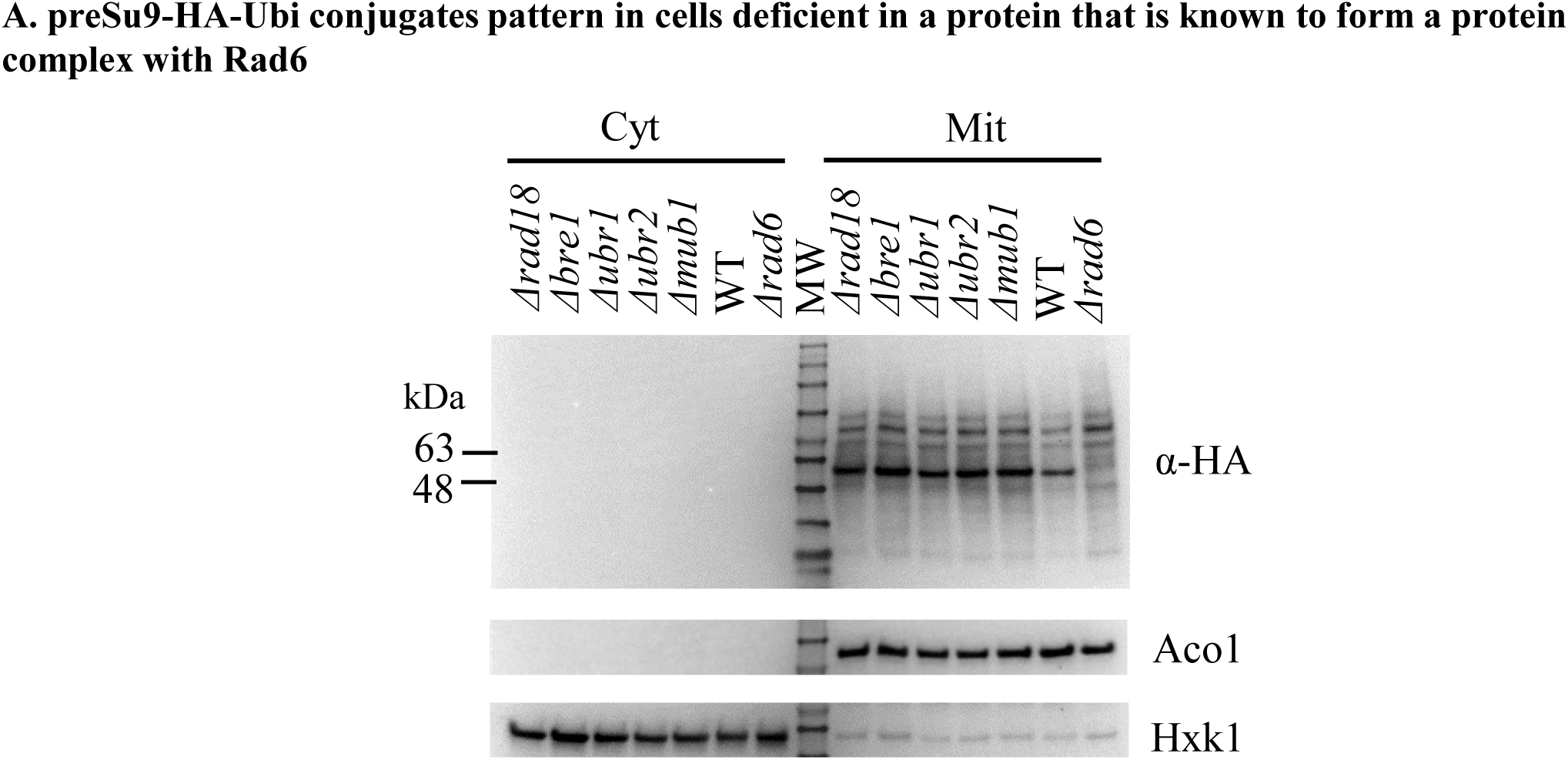
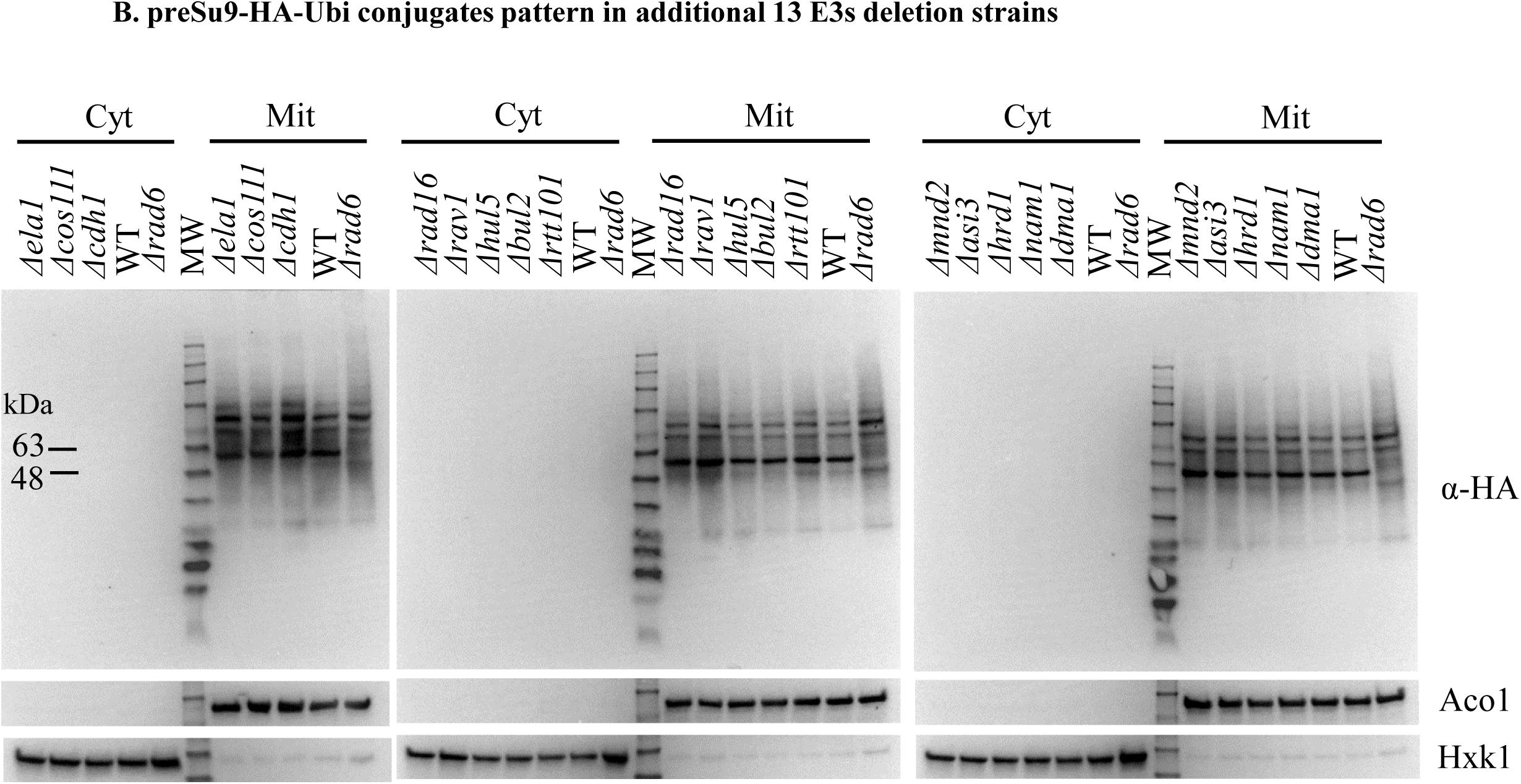
preSu9-HA-Ubi conjugate patterns in cells deficient in E3 (related) proteins. **(A)** Rad18, Bre1, Ubr1, Ubr2, and Mub1 are the 5 known proteins that form complexes with Rad6 in ubiquitinating substrate proteins. Cytosolic and mitochondrial fractions of indicated cells expressing preSu9-HA-Ubi were examined by WB with antibody against HA, Aco1 (mitochondrial marker), and Hxk1 (cytosolic marker). **(B)** Cytosolic and mitochondrial fractions of cells deficient in an indicated E3s and expressing preSu9-HA-Ubi were examined by WB with antibody against HA, Aco1 (mitochondrial marker), and Hxk1 (cytosolic marker).

**Figure S5:**
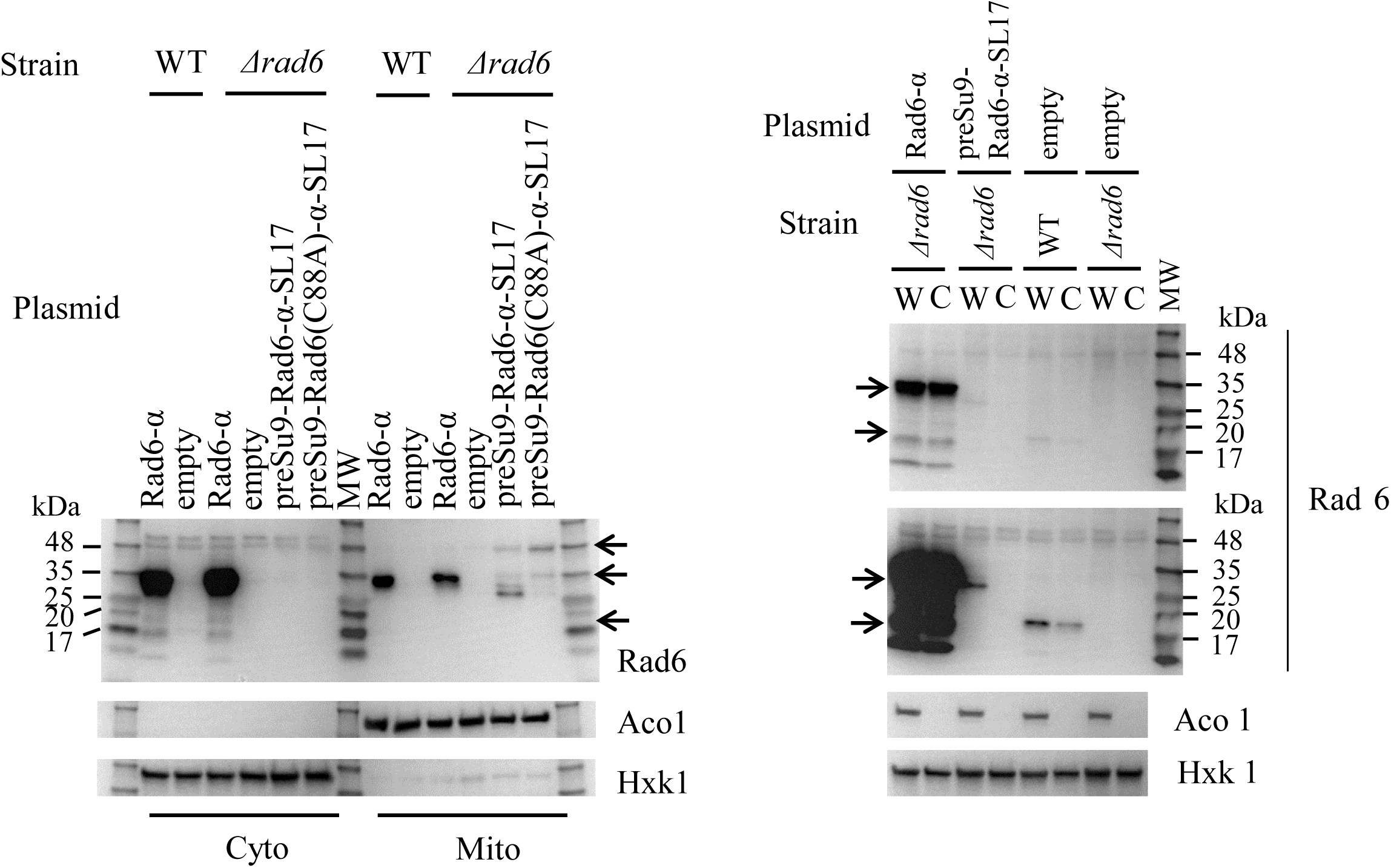
**Native level of Rad6 is low** Cytosolic (Cyto or C), mitochondrial fractions (Mito) and whole cell extractions (W) prepared from WT or *Δrad6* cells harboring empty plasmid or a Rad6 construct were examined by WB with antibody against Rad6, Aco1 (mitochondrial marker), and Hxk1 (cytosolic marker). Arrows indicate the migrating position of Rad6 or Rad6 fusion proteins. The 1^st^ and 2^nd^ panels of the right picture are produced respectively by shorter and longer exposure for the probing with Rad6 antibody.

**Figure S6.**
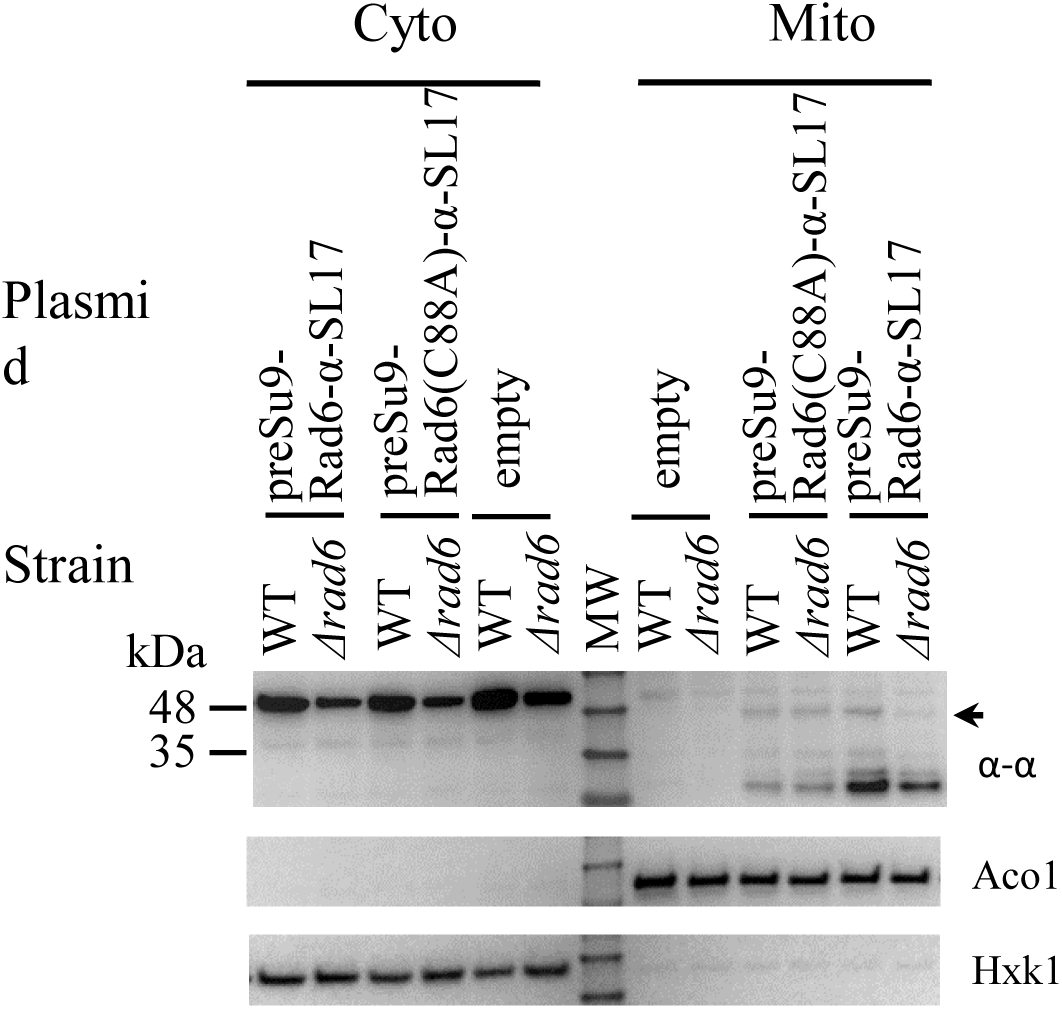
(for. **Figure 7E) Expression of preSu9-Rad6(C88A)-α-SL17.** Subcellular fractions of WT or *Δrad6* cells harbouring the indicated plasmids were probed for the α-tag signal using WB. The arrow indicates the full-length Rad6 fusion protein. Aco1, mitochondrial marker; Hxk1, cytosolic marker.

**Table S2:**
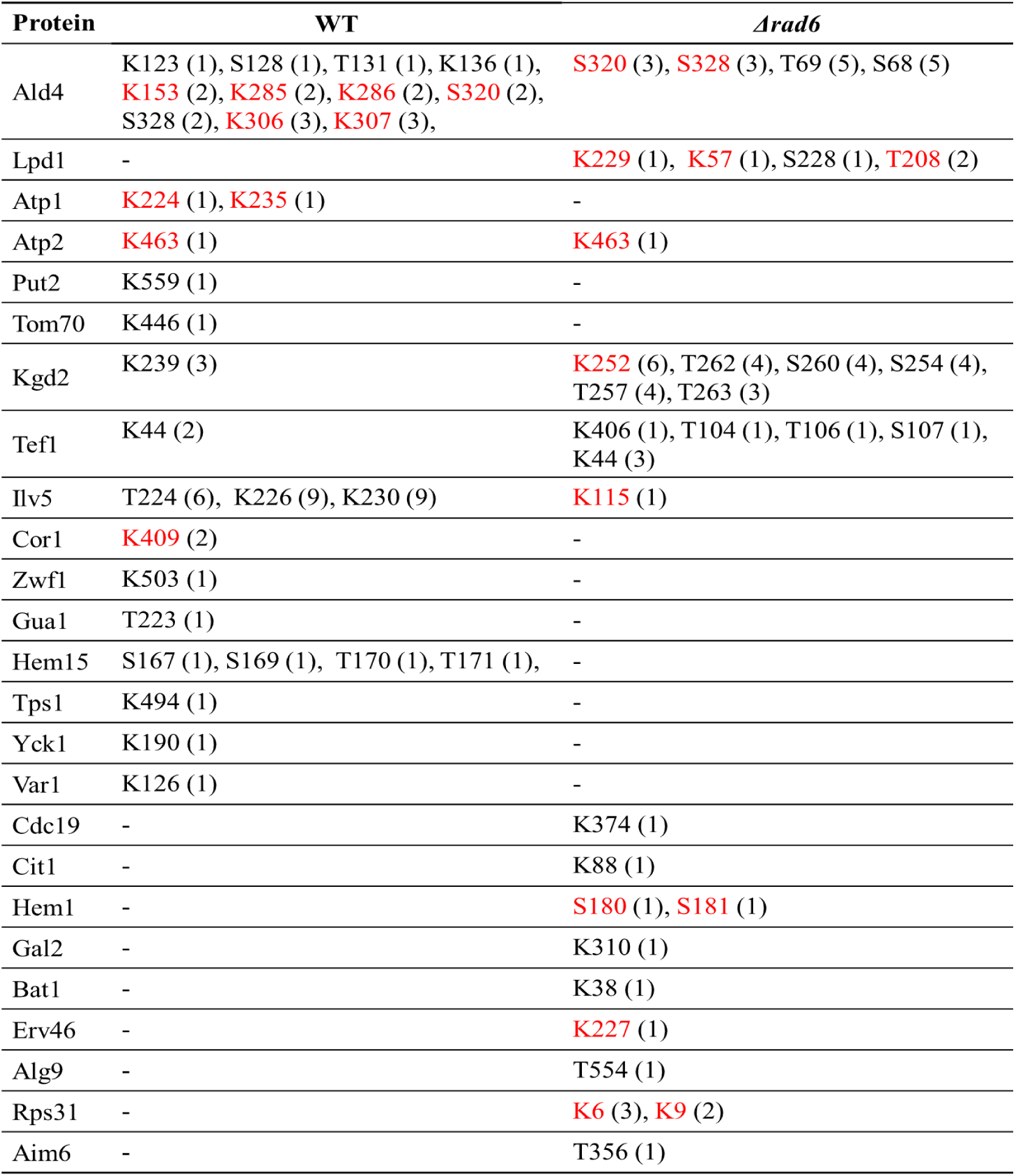
Ubiquitin modifications detected. Mitochondrial lysates of WT and *Δrad6* cells expressing preSu9-HA-Ubi were subjected to SDS-PAGE followed by gel staining with InstantBlue. The proteins separated between 48 and 63 kDa were excised for each sample and subjected to MS. The detected ubiquitin modifications are summarized. Modifications with case(s) with a confidence level higher than 90% are highlighted in red. The number in the brackets indicate the number of corresponding modifications observed by MS.

**Figure S7:**
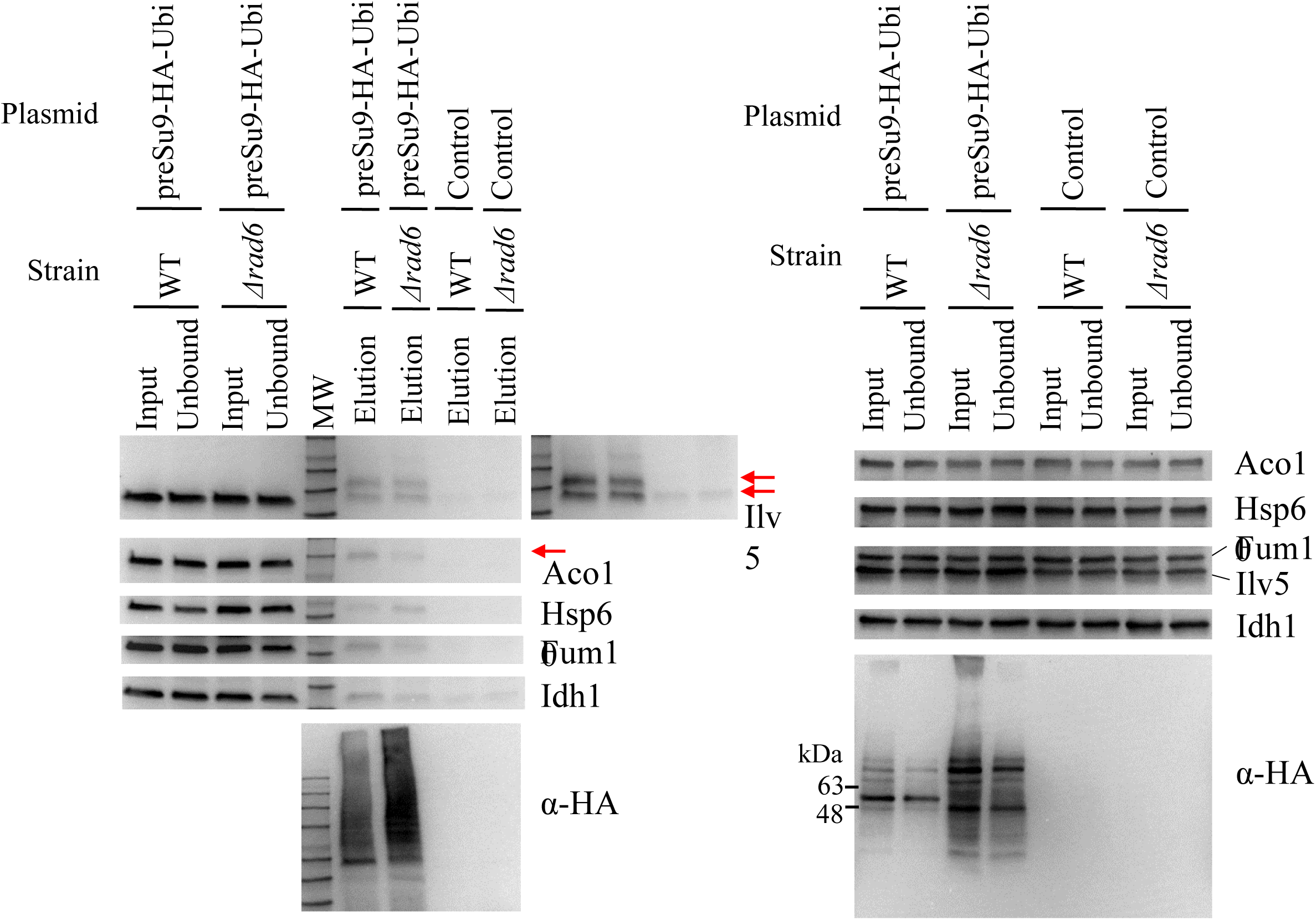
preSu9-HA-Ubi IP from mitochondrial lysates of WT versus *Δrad6* cells. Left panel: IPs with anti-HA magnetic beads incubated with mitochondrial lysates (containing ∼1 mg of proteins) of cells harboring empty vector (Control) or preSu9- HA-Ubi. Proteins were eluted from immunoprecipitated beads by boiling the beads in protein loading dye. Input and unbound samples, and 1/5 of elution samples were analyzed by western blot. A longer time exposure of anti-Ilv5 antibodies is shown on the right. Red arrows indicate the protein band of the estimated ubiquitinated form. Right panel: Input and unbound samples of IPs analyzed by WB.

